# IRAK4 autophosphorylation controls inflammatory signaling by activating IRAK oligomerization

**DOI:** 10.1101/2023.12.21.572799

**Authors:** Niranjan Srikanth, Rafael Deliz-Aguirre, Deepika Kumari Gola, Margaux Bilay, Elke Ziska, Marcus J. Taylor

## Abstract

The controlled oligomerization of signaling proteins is an essential feature of many inflammatory signaling pathways. An example is IL-1 receptor signaling, which relies on the oligomerization of the Death Domain (DD)-containing proteins MyD88 and IRAK family kinases. This process leads to the assembly of the Myddosome signaling complex, and disrupting assembly holds potential for anti-inflammatory treatments. However, IRAKs’ signaling activity is also regulated by auto-/trans-phosphorylation, and it is unclear if these processes operate at or downstream of Myddosome assembly. Here, we find that the initial stage of Myddosome assembly is solely controlled by MyD88:IRAK4 DD interactions. In later stages, IRAK4 auto-phosphorylation serves as a switch, regulating IRAK1/2/3 incorporation and DD oligomerization. Small molecule inhibitors of IRAK4 kinase activity block this later stage of assembly, explaining how they dampen inflammatory signaling. Our data reveals IRAK4 auto-phosphorylation as an energy-dependent switch activating the heterotypic assembly of IRAKs’ DDs and downstream inflammatory IL-1 signaling. This highlights how a signaling cascade integrates phosphorylation and protein oligomerization steps.

## Introduction

Activation of Interleukin 1 receptors (IL-1R) is essential for the innate immune response, but dysregulated signaling from these receptors can cause inflammatory disorders. A key player in regulating IL-1R signal transduction is the Myddosome, an oligomeric immune complex^1,2^. Myddosome formation requires the recruitment of MyD88 to the activated receptor, and the controlled coassembly with IRAK4 and either IRAK1, IRAK2 or IRAK3. The core of this complex is a mixed oligomer of MyD88 and IRAK Death domains (DD)^2,3^. Therefore an essential step in IL-1R signal transduction is controlled DD oligomerization and IRAK co-assembly with MyD88. In addition to a n-terminus DD, IRAKs also have a c-terminus kinase domain^4^, which presents a possible therapeutic target for treating inflammatory disorders. However, the IRAK family contains possible pseudokinases as well as active kinases^5–8^, and genetic studies that have suggested that IRAKs have both redundant and distinct functional roles in signal transduction^9–11^. Therefore, the functional relationship of IRAK DD oligomerization and the kinase domain remains poorly understood.

Of all the IRAKs, IRAK4 is a non-redundant and essential Myddosome component^1,2,12^. Beyond a structural role of the DD in Myddosome formation, IRAK4 can auto-phosphorylate and phosphorylate other IRAK family members^8,13^. The IRAK4 kinase activity is required for downstream signaling^14–16^, making it an attractive therapeutic target. However when and where in the signaling pathway the IRAK4 kinase activity operates remains unknown. A critical question is what is the regulatory relationship between DD self-assembly in Myddosome formation, IRAK4 autophosphorylation and downstream phosphorylation of other IRAKs. Resolving these discrete processes and the molecular logic of how Myddosome assembly is regulated and controlled to transduce signals is essential to understand how cells activate innate immune responses.

Here we analyzed the role of IRAK4 kinase activity and autophosphorylation in IL-1 signaling transduction. We found that auto-phosphorylation of the IRAK4 kinase domain is a discrete stage of Myddosome assembly, and occurs after IRAK4 and MyD88 co-assembly. Using live cell microscopy to capture the dynamics of myddosome formation, we observed that pharmacological inhibition of IRAK4 kinase activity or kinase inactive mutants still resulted in co-assembly with MyD88. This finding indicates that death domain interactions are the sole requirement for MyD88:IRAK4 co-assembly. Instead, IRAK4 auto-phosphorylation triggers the stable recruitment of IRAK1. This is likely mediated by heterodimerization of the phosphorylated IRAK4 kinase domain and the IRAK1 kinase domain, which triggers the incorporation and oligomerization of the IRAK1 death domain at nascent Myddosomes. This regulatory mechanism also controlled the incorporation of IRAK2 and IRAK3 into Myddosome complexes. Our data suggests that IRAK4 autophosphorylation is a discrete intermediate stage of myddosome assembly. This energy dependent step functions as a regulatory switch that controls the final stages of DD assembly in Myddosome formation. Our work highlights how protein oligomerization can be regulated by enzymatic activity within a signaling cascade.

## Results

### The co-assembly of IRAK4 with MyD88 and IRAK1 is coincident with IRAK4 autophosphorylation

Myddosomes form by a sequential choreography of DD interaction starting with MyD88 multimerization which triggers the recruitment of IRAK4 and finally IRAK1^2,17^. IRAK4 death domain self-assembly and kinase activity are essential to Myddosome signaling^14–16,18^. A critical question is if and at what point IRAK4 kinase activity and phosphorylation regulates this macromolecular assembly reaction. To dissect the stages of Myddosome formation, we used gene-edited EL4 cells expressing MyD88-GFP/IRAK4-mScarlet or MyD88-GFP/IRAK1-mScarlet. Employing IL-1 functionalized supported lipid bilayers (SLBs), we activated cells and visualized Myddosome formation via total internal reflection (TIRF) microscopy ^17,19^. Kymograph analysis shows nucleation of MyD88 puncta followed by subsequent recruitment of either IRAK4 and IRAK1 (Fig. 1A, C). We observe that the number of MyD88 co-assemblies with IRAK4 or IRAK1 continuously increases over 18 minutes within single live EL4 cells (Fig. 1B, D, Movie S1 and S2). We conclude that MyD88 coassembly with IRAK4 and IRAK1 is triggerable, and increases in a time dependent manner post IL-1 stimulation.

**Figure 1.**
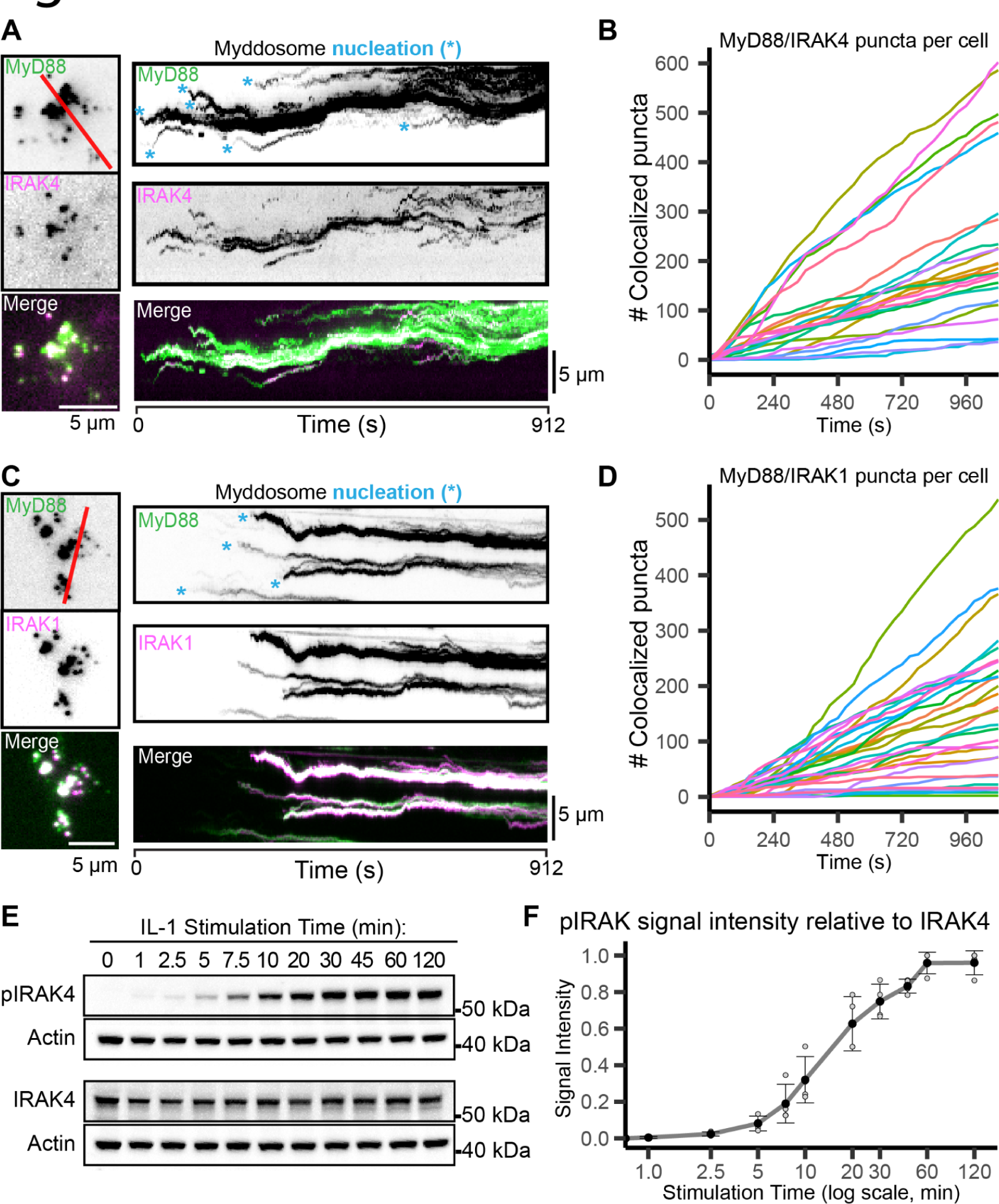
Time course analysis of Myddosome formation and pIRAK4 production is consistent with IRAK4 kinase activity involvement in Myddosome assembly **A)** TIRF images of EL4-MyD88-GFP/IRAK4-mScarlet stimulated on IL-1 functionalized SLBs. Kymographs derived from red line overlaid TIRF images (left panel). Scale bar, 5 µm. The blue asterisks on the MyD88-GFP kymograph highlight the nucleation of Myddosome complexes. **B)** Formation of MyD88-GFP:IRAK4-mScarlet assemblies over time for individual cells (each color represents one of 24 cells). t = 0 is defined as the time point when the cell lands on the bilayer. **C)** TIRF images of EL4-MyD88-GFP/IRAK1-mScarlet stimulated on IL-1 functionalized SLBs. Kymographs derived from red line overlaid TIRF images (left panel). Scale bar, 5 µm. The blue asterisks on the MyD88-GFP kymograph highlight the nucleation of Myddosome complexes. **D)** Formation of MyD88-GFP:IRAK1-mScarlet assemblies over time for individual cells (each color represents one of 31 cells). t = 0 is defined as the time point when the cell lands on the bilayer. **E)** Time course Western blot analysis of pIRAK4 levels in EL4 cells after IL-1 stimulation (1 ng/ml). Lysates were separately probed for antibodies specific for pIRAK4 (Thr-345/Ser-346) and total IRAK4. Actin detection was used as a loading control to normalize the band intensity. **F)** Time course quantification of the pIRAK4 signal intensity relative to total IRAK4 signal intensity from the Western blots shown in (e). Each black colored data point is the average relative pIRAK level derived from N=4 experimental replicates (see Supplementary Fig. 1A). Smaller data points are the relative pIRAK signal intensity of individual replicates.

IRAK4 auto phosphorylates serine/threonine residues in its kinase domain^8,20^. This reaction occurs between dimerized kinase domains (trans-autophosphorylation), and the presence of MyD88 increases the rate of this reaction *in vitro*^13^. This suggests that IRAK4 multimerization during Myddosome assembly could stimulate auto-phosphorylation, and the time course of phosphorylated-IRAK4 (pIRAK4) production would be matched to the rate of Myddosome complex formation (Fig. 1B, D). We analyzed the time course of IRAK4 auto-phosphorylation using Western blot analysis with an antibody that recognizes the phosphorylated Thr-345/Ser-346 residues of IRAK4^21^ (Fig. 1E, Supplementary Fig. 1A). This analysis reveals that phosphorylation of IRAK4 occurs post IL-1 stimulation and the level of pIRAK4 increases over the first 20-30 mins post IL-1 stimulation (Fig. 1F). The kinetics of pIRAK4 production in EL4 cells is comparable to what has been reported across multiple cell lines^22,23^, and when we observe Myddosome assembly (Fig. 1A, C). This demonstrates that Myddosome assembly and IRAK4 phosphorylation are dynamic processes, and that the number of Myddosome complexes and pIRAK4 levels increases in a time dependent manner post IL-1 stimulation. From this data we conclude that phosphorylation of IRAK4 coincides with DD assembly and myddosome formation. This is consistent with IRAK4 auto-phosphorylation occurring as IRAK4 co-assembles with MyD88 and IRAK1.

### IRAK4 autophosphorylation is a discrete step in myddosome assembly

To determine if IRAK4 auto-phosphorylation occurs within Myddosomes and regulates assembly, we visualized the subcellular localization of pIRAK4 and Myddosome components using EL4 knock-in cells expressing MyD88-GFP and either IRAK4-mScarlet or IRAK1-mScarlet cells^17^. We stimulated cells for 15 mins with IL-1 functionalized SLBs, and then fixed and immuno-labeled cells with a pIRAK4 specific antibody^21^. In this system, Myddosomes assemble asynchronously as EL4 cells come into contact with the IL-1 functionalized SLB at different times. Myddosome complexes continue to nucleate and form over 15 minutes within individual cells (Fig. 1B, D). Consequently, chemical fixation would capture Myddosomes at various assembly stages. If IRAK4 autophosphorylation is triggered at a specific stage or regulates Myddosome assembly, we anticipate observing partially assembled Myddosomes both with and without pIRAK4. These would colocalize with specific combinations of MyD88, total IRAK4, or IRAK1. In MyD88-GFP/IRAK4-mScarlet expressing cells, we observed MyD88 puncta with IRAK4-mScarlet, both with and without pIRAK4 (Fig. 2A, B). Staining of MyD88-GFP/IRAK1-mScarlet cells revealed puncta composed of MyD88, MyD88 with pIRAK4, and MyD88 with pIRAK4 and IRAK1 (Fig. 2C, D). More than 80% of MyD88 puncta with IRAK1 contained pIRAK4. These combinations likely represent fully and partially assembled Myddosomes. Considering the distribution of pIRAK4 staining, and that IRAK4 phosphorylation is a post-stimulation event (Fig. 1E, Fig. 2B, D), we conclude that IRAK4 is likely recruited to the MyD88 oligomer in an unphosphorylated state, and its assembly with MyD88 triggers auto-phosphorylation.

**Figure 2:**
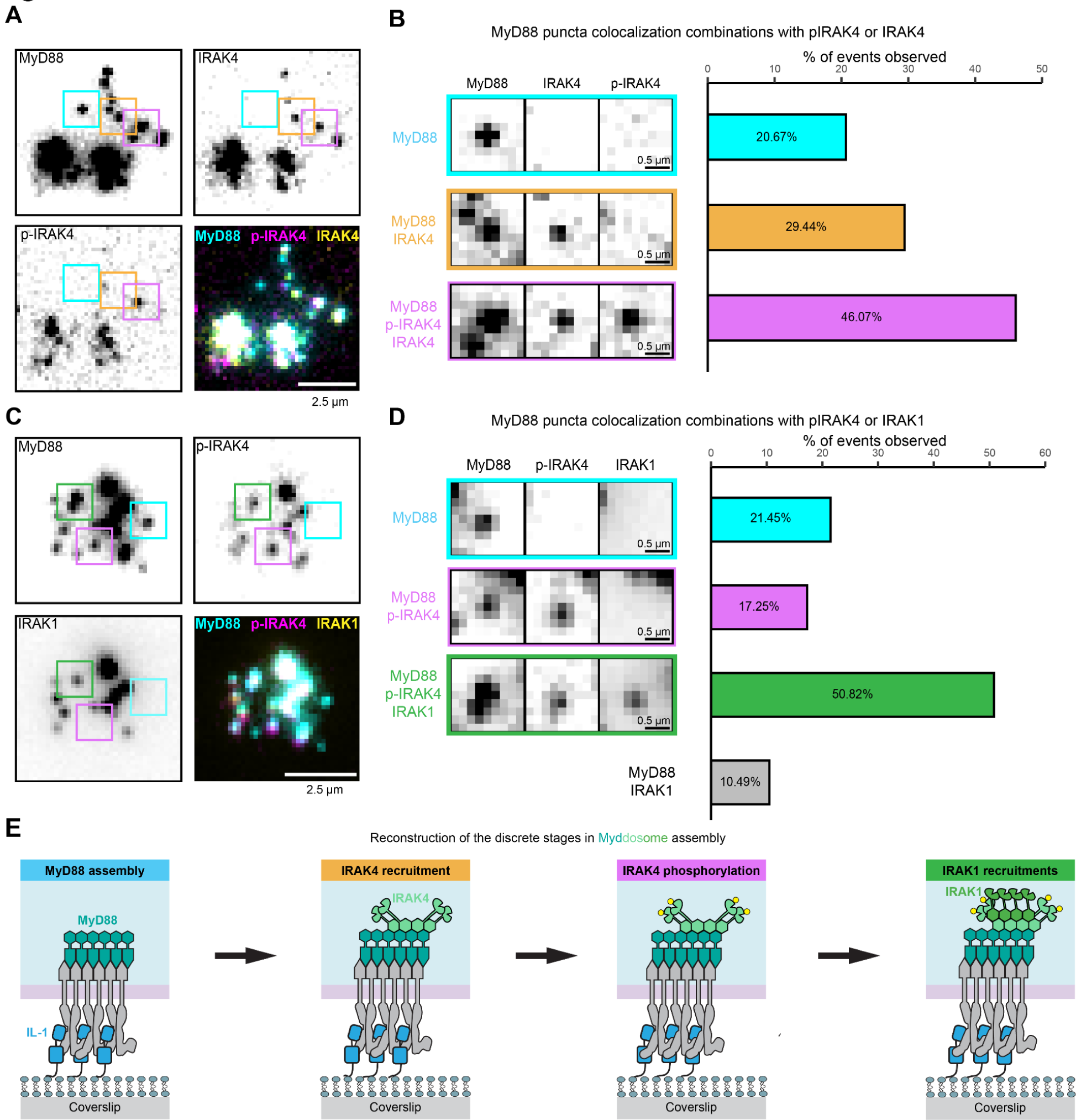
pIRAK4 localizes to Myddosome suggesting that IRAK4 autophosphorylation is a discrete stage of Myddosome assembly **a)** TIRF images of fixed EL4-MyD88-GFP/IRAK4-mScarlet cells stained with antibodies against pIRAK4. Bottom right panel is a merge of all channels. Cells were activated on IL-1 functionalized SLBs for 20 min before fixation. Coloured boxes highlight assemblies that either only contain MyD88 (cyan), MyD88 and IRAK4 (yellow-orange) and MyD88, IRAK4 and pIRAK4 (magenta). Scale bar, 2.5 µm. **b)** Image panel (left) shows the highlighted boxes in (a) comparing the same spot regions in different channels. Bar plot (right) shows percentage of MyD88 puncta that colocalize with IRAK4 or pIRAK4 and IRAK4. Bar chart generated from 445 MyD88 puncta. **c)** TIRF images of fixed EL4-MyD88-GFP/IRAK1-mScarlet cells stained with antibodies against pIRAK4. Bottom right panel is a merge of all the channels. Cells were activated on IL-1 functionalized SLBs for 20 min before fixation. Coloured boxes highlight MyD88 assemblies that either only contain MyD88 (cyan), MyD88 and pIRAK4 (magenta), MyD88, pIRAK4 and IRAK1 (green). Scale bar, 2.5 µm. **d)** Image panel (left) shows the highlighted boxes in (a) comparing the same spot regions in different channels. Bar plot (right) shows percentage of MyD88 puncta that colocalize with pIRAK4, pIRAK4 and IRAK1 or IRAK1. Bar chart generated from 429 MyD88 puncta. **e)** A reconstructed timeline showing the stages of Myddosome assembly interpreted from the pIRAK4 staining patterns in cells expressing MyD88-GFP and IRAK4-mScarlet (a) or MyD88-GFP and IRAK1-mScarlet (c).

Based on the relative proportion of MyD88 puncta colocalized with IRAK4, pIRAK4 and IRAK1 (Fig. 2B, D), we reconstructed a timeline of events where phosphorylation of IRAK4 occurs post IRAK4 recruitment and pre IRAK1 recruitment (Fig. 2E). This suggests that the self-assembly of DDs can be regulated by the energy dependent step of IRAK4 autophosphorylation. Thus kinase activity may not be strictly required for IRAK4 recruitment, but could regulate the stability of MyD88:IRAK4 complex or IRAK1 recruitment. This is consistent with the kinase activity and IRAK4 auto-phosphorylation functioning as a regulatory switch in Myddosome assembly and downstream signal transduction.

### Pharmacological inhibition of IRAK4 kinase activity does not interfere with MyD88:IRAK4 co-assembly

If IRAK4 kinase activity acts upon DD assembly and therefore regulates Myddosome formation, inhibition of IRAK4 kinase activity and auto-phosphorylation should disrupt assembly or freeze the process at a specific stage (Fig. 2E). One of the initial stages of formation is the co-assembly of MyD88 and IRAK4 mediated by the heterotypic interactions between the respective DDs (Fig. 2E)^2,3^. We assayed the dynamics of MyD88 and IRAK4 co assembly in live cells in the presence of a IRAK4 kinase inhibitor^24^, that blocks downstream signaling and inhibited IRAK4 autophosphorylation (Supplementary Fig. 2A, B). In the presence of this inhibitor we observe the formation of MyD88 assemblies that recruited IRAK4 (Fig. 3A, Supplementary Fig. 2C, Movie S3). We found no differences in the percentage of IRAK4 colocalized MyD88 puncta (Fig. 3B). We analyzed the dynamics and properties of individual MyD88:IRAK4 assemblies (Fig. 3C), which in both control and inhibitor treated cells showed the nucleation and growth of MyD88 preceded the recruitment of IRAK4. IRAK4 has been shown to control MyD88 oligomerization^17^. If kinase activity of IRAK4 negatively regulated MyD88 oligomerization, we would expect larger MyD88 assemblies. However, the size of MyD88 assemblies was identical in control and inhibitor treated cells (Fig. 3D). We conclude that IRAK4 kinase activity does not affect the initial stage of MyD88 oligomerization.

**Figure 3:**
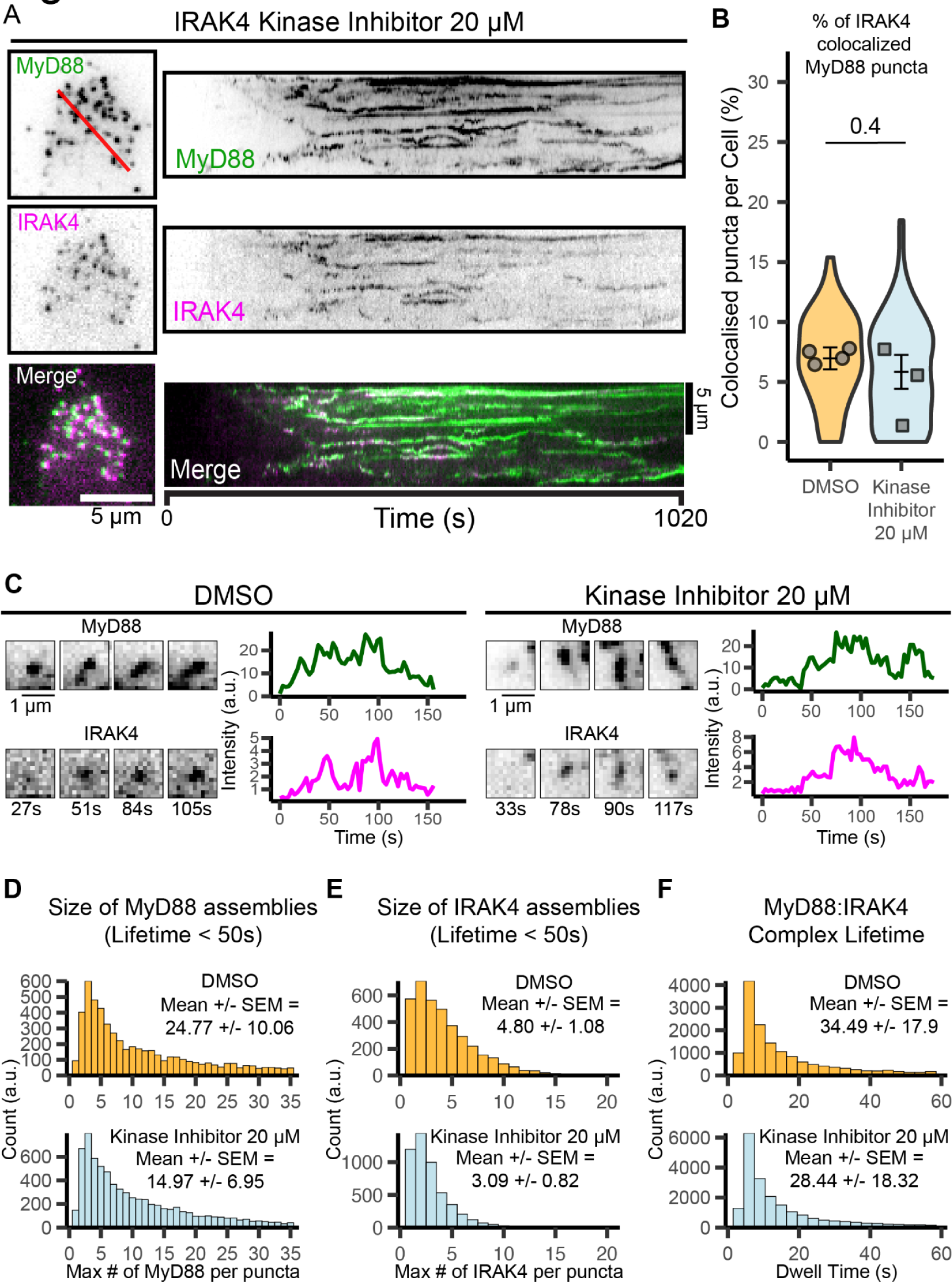
Pharmacological inhibition of IRAK4 kinase activity does not perturb IRAK4:MyD88 co-assembly **a)** TIRF images of EL4-MyD88-GFP/IRAK4-mScarlet treated with IRAK4 kinase inhibitor stimulated on IL-1 functionalized SLBs. Kymographs derived from red line overlaid TIRF images (left panel). Scale bar, 5 µm. **b)** Quantification of the % of MyD88-GFP puncta that colocalize with IRAK4-mScarlet puncta. Violin plots show the distribution of the cell data. Data points superimposed on violin plots are the averages of independent experimental replicates. Bars represent mean ± SEM (n = 4 and 3 experimental replicates, each replicate encompasses measurements from 3-32 and 17-46 cells, from the DMSO and inhibitor treatment conditions respectively). Statistical significance is determined using the Wilcoxon Rank-Sum test. **c)** Time-series TIRF images and fluorescence-intensity time series showing the formation and coassembly of MyD88-GFP and IRAK4-mScarlet puncta from cells treated with DMSO (left panel) or IRAK4 kinase inhibitor (right panel). Scale bar, 1 µm. **d-e**) Histogram showing the distribution of the size of MyD88 (d) and IRAK4 (e) assemblies of DMSO (top, yellow) and IRAK4 kinase inhibitor (bottom, light blue) treated cells; n = 6568 versus n = 7823 MyD88 puncta from the DMSO and inhibitor treatment conditions respectively, MyD88 puncta pooled from 4 experimental replicates (d); n = 3869 versus n = 5067 IRAK4 puncta from the DMSO and inhibitor treatment conditions, IRAK4 puncta pooled from 3 experimental replicates (e). **f)** Distribution of the lifetime of the MyD88:IRAK4 complex in DMSO and IRAK4 kinase inhibitor treated cells. Mean ± SEM derived from 4 and 3 experimental replicates for each condition. n = 16732 versus n = 22652 MyD88:IRAK4 assemblies from DMSO and inhibitor treatment conditions, pooled across experimental replicates.

IRAK4 auto-phosphorylation might regulate the stability of the MyD88:IRAK4 complex or enhance IRAK4 oligomerization at assembling Myddosomes. We analyzed the lifetime of MyD88:IRAK4 complexes, and found the distribution was equivalent in both control and inhibitor treated cells (Fig. 3F). We found that the size of IRAK4 assemblies was equivalent (Fig. 3E), suggesting inhibition of the kinase activity does not perturb IRAK4 oligomerization. This suggests that IRAK4 kinase activity does not regulate the formation of the MyD88:IRAK4 complex, and auto-phosphorylation does not participate in the initial two stages of Myddosome formation (Fig. 2E). These signal transduction steps are likely mediated by homotypic and heterotypic interactions between the MyD88 and IRAK4 DDs^2,3^. We conclude that IRAK4 auto-phosphorylation and kinase activity must regulate signaling reactions downstream of these stages.

### MyD88:IRAK4 assembly is solely controlled by Death domain interactions

The unperturbed dynamics of MyD88:IRAK4 co-assembly in the presence of kinase inhibition, suggest that auto-phosphorylation does not regulate the initial stages of Myddosome assembly. This implies that heterotypic and homotypic interactions between the MyD88 and IRAK4 DD solely regulate MyD88:IRAK4 co-assembly and IRAK4 multimerization. To confirm whether these initial stages are controlled by DD multimerization, we used a genetic reconstitution approach. We generated a kinase inactive version of IRAK4 (IRAK4^K213/214A^) and a truncated IRAK4 solely containing the DD (IRAK4^DD^) tagged with mScarlet (Fig. 4A). These constructs along with full length wild type (WT) IRAK4 (IRAK4^WT^), were reconstituted into EL4-MyD88-GFP/IRAK1-KO cells^15^. We found IRAK4^K213/214A^ and IRAK4^DD^ could not restore IL-1 signaling responses in IRAK4 KO cells (Supplementary Fig. 3A), consistent with the kinase domain and activity being critical for signal transduction. However, we found that MyD88 can co-assemble with both the IRAK4^K213/214A^ and IRAK4^DD^ (Fig. 4B, C, Supplementary Fig. 3B, Movie S4), and the percentage of colocalized assemblies was equivalent to KO cells reconstituted with IRAK4^WT^ (Fig. 4D). Thus MyD88:IRAK4 co-assembly is solely dependent on DD:DD interactions, and is not regulated by the IRAK4 kinase domain or activity. Analysis of individual MyD88 puncta found that they recruited and co-assembled with both IRAK4^K213/214A^ and IRAK4^DD^ with similar kinetics (Fig. 4E, Supplementary Fig. 3C) and expression of both mutant alleles had no effect on the size of MyD88-GFP assemblies (Fig. 4F). WT and mutant IRAK4 alleles formed assemblies with equivalent size distributions (Fig. 4G), suggesting that IRAK4 multimerization was solely controlled by DD interactions. Finally, co-assemblies of MyD88 with WT or mutant IRAK4 alleles had equivalent lifetime distributions (Fig. 4H), suggesting the IRAK4 kinase domain and activity does not impact the stability of the MyD88:IRAK4 complex. We conclude that the initial steps of signal transduction and Myddosome formation are regulated by DD assembly, with no energy dependent post-translational modifications from the IRAK4 kinase activity. If the IRAK4 kinase domain and auto-phosphorylation are part of a regulatory switch controlling signal transduction, this switch would operate downstream of MyD88:IRAK4 co-assembly and multimerization.

**Figure 4.**
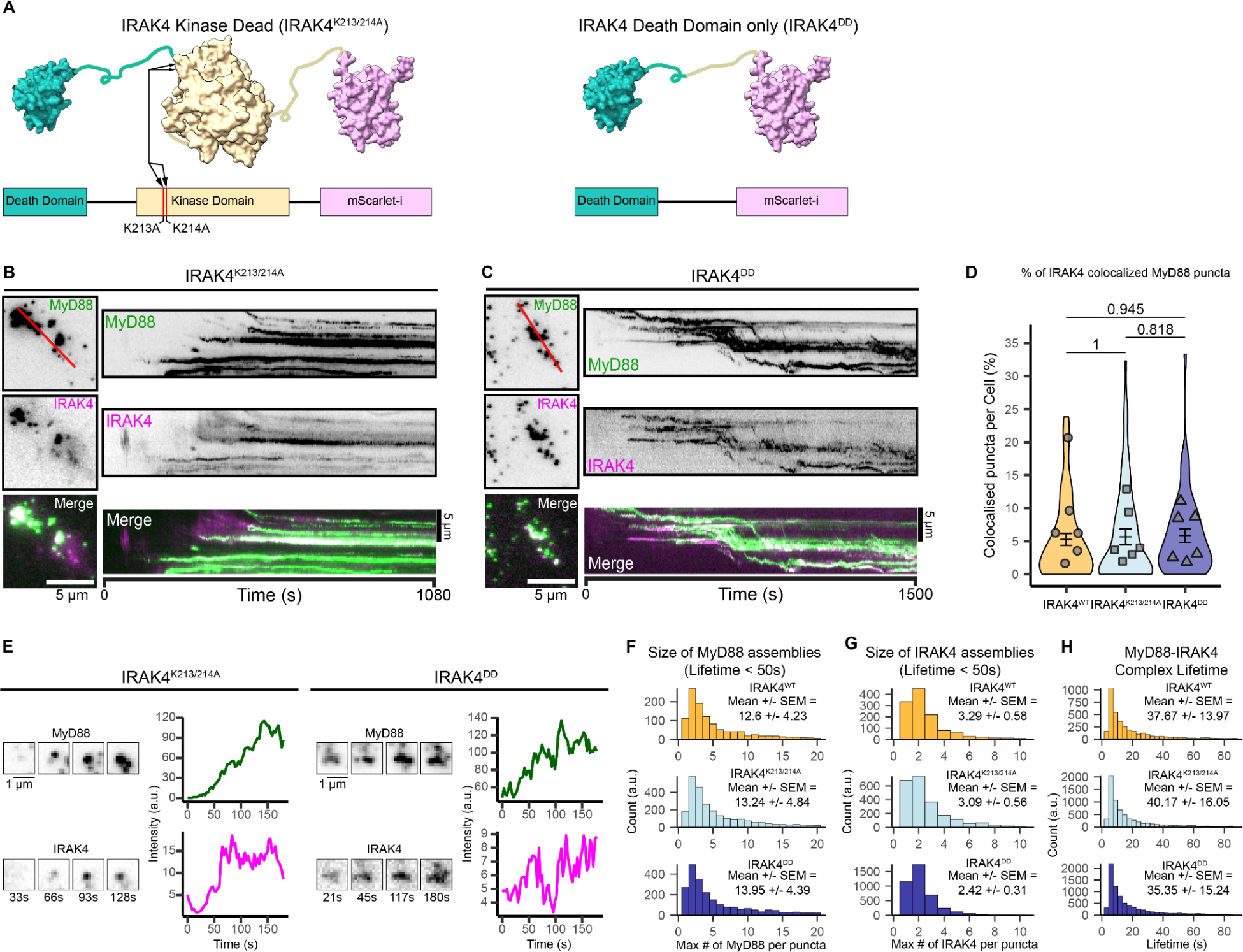
IRAK4 kinase activity and kinase domain are dispensable for the assembly of an oligomeric MyD88:IRAK4 complex **a)** Domain schematic of mutant IRAK4 alleles tagged with mScarlet and reconstituted into EL4-MyD88-GFP/IRAK4-KO. **b-c**) TIRF images of EL4-MyD88-GFP/IRAK4-KO expressing IRAK4^K213/214A^ (b) or IRAK4^DD^ (c) stimulated on IL-1 functionalized SLBs. Kymographs derived from red line overlaid TIRF images (left panel). Scale bar, 5 µm. **d)** Percentage of IRAK4 colocalization with MyD88 in EL4-MyD88-GFP/IRAK4-KO cells expressing IRAK4^WT^, IRAK4^K213/214A^ and IRAK4^DD^ cells. IRAK4^WT^, IRAK4^K213/214A^ and IRAK4^DD^. Violin plots show the distribution of individual cell measurements. The dots superimposed on the violin plots represent the mean value of each replicate (n = 6, 6 and 6 replicates with each replicate encompassing measurements from 2-15, 8-23, 5-14 cells for IRAK4^WT^, IRAK4^K213/214A^ and IRAK4^DD^ respectively). **e)** Time-series TIRF images and fluorescence-intensity time series showing the formation of a single MyD88-GFP:IRAK4^K213/214A^ (left panel) or MyD88-GFP:IRAK4^DD^ assembly (right panel) . Scale bar, 1 µm. **f-g**) Histogram plots showing the size distribution of (f) MyD88 assemblies and (g) IRAK4 assemblies in EL4-MyD88-GFP/IRAK4-KO cells reconstituted with IRAK4^WT^ (upper, yellow), IRAK4^K213/214A^ (middle, light blue) and IRAK4^DD^ (lower, dark blue). In panel (f), histograms are composed of n = 465, 2778 and 2424 MyD88 assemblies from cells expressing IRAK4^WT^, IRAK4^K213/214A^ and IRAK4^DD^ respectively. In panel (g), histograms are composed of n = 605, 1958 and 3857 IRAK4 assemblies from cells expressing IRAK4^WT^, IRAK4^K213/214A^ and IRAK4^DD^ respectively. **h**) Histogram plots of MyD88:IRAK4 complex lifetime distribution in EL4-MyD88-GFP/IRAK4-KO cells reconstituted with IRAK4^WT^ (upper, yellow), IRAK4^K213/214A^ (middle, light blue) and IRAK4^DD^ (lower, dark blue). Histograms are composed of n = 1787, 7098 and 7386 MyD88:IRAK4 assemblies from cells expressing IRAK4^WT^, IRAK4^K213/214A^ and IRAK4^DD^ respectively.

### Pharmacological inhibition of IRAK4 kinase activity reduces the stable incorporation of IRAK1 into nascent Myddosomes

The final assembly stage of Myddosome complexes is the recruitment of IRAK1 to MyD88:IRAK4 assemblies. Our timeline reconstruction shows that IRAK1 is recruited onto complexes that possess pIRAK4 (Fig. 2E). This indicates that phosphorylation of IRAK4 may be required for IRAK1 recruitment. We tested this by treating the EL4-MyD88-GFP/IRAK1-mScarlet cell line with the inhibitor. We observed less MyD88-IRAK1 assemblies in inhibitor treated cells (Fig. 5A, B, Movie S5). In addition, IRAK1-mScarlet puncta appeared to have decreased intensity and a shorter dwell time. We observed that treatment with the IRAK4 kinase inhibitor decreased the percentage of MyD88 puncta that colocalized with IRAK1 (Fig. 5C). We observed the maximal effect at 20 µM IRAK4 kinase inhibitor which was 3 fold lower percentage of IRAK1 colocalized MyD88 puncta (2% and 6.9% for kinase inhibitor versus DMSO treated cells respectively, Fig. 5C). In addition to frequency of IRAK1 recruitment, we analyzed whether IRAK4 kinase inhibition changed the properties of IRAK1 assemblies. At both 500 nM and 20 µM the size of IRAK1 assemblies that colocalized with MyD88 puncta was reduced (Fig. 5D). We found that treatment with IRAK4 kinase inhibitor reduced the lifetime of IRAK1 colocalized MyD88 puncta (Fig. 5E), suggesting the MyD88:IRAK4:IRAK1 complex had reduced stability when IRAK4 kinase activity was inhibited. We conclude that IRAK4 kinase activity and auto-phosphorylation regulates IRAK1 recruitment and stable incorporation into Myddosomes.

**Figure 5:**
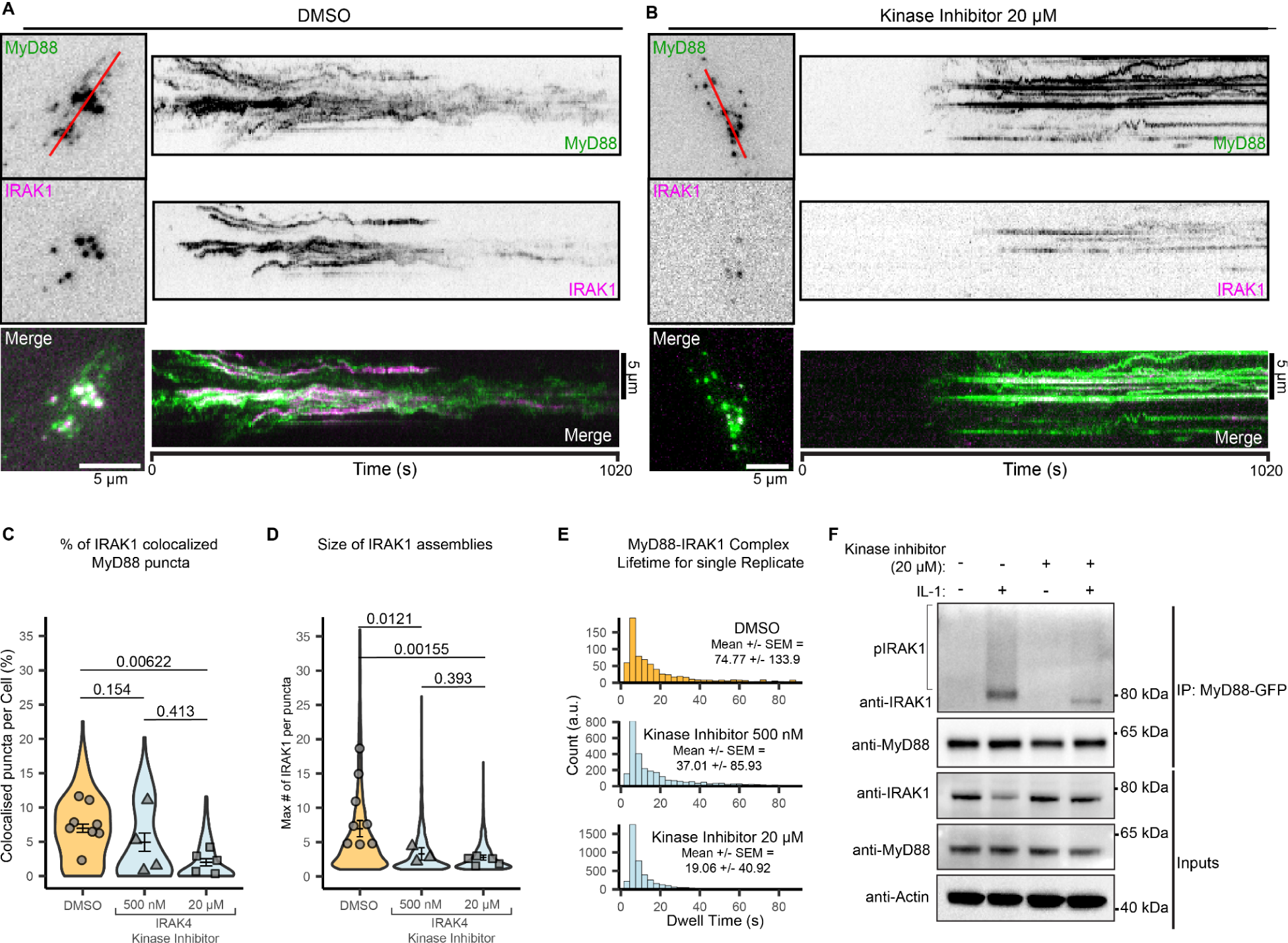
Pharmacological inhibition of IRAK4 Kinase activity stops IRAK1 recruitment and incorporation into nascent Myddosomes **a-b**) Kymographs of DMSO control (a) or IRAK4 kinase inhibitor (b) treated EL4-MyD88-GFP/IRAK1-mScarlet cells. Kymographs derived from red line overlaid TIRF images (left panel). Scale bar, 5 µm. **c)** Percentage of MyD88 puncta that colocalize with IRAK1 puncta per cell. Violin plots show the distribution of individual cell measurements for DMSO, 500 nM and 20 µM kinase inhibitor treatment. The dots superimposed on the violin plots represent the mean value of each replicate (n = 8, 4 and 5 replicates with each replicate encompassing measurements from 12-32, 15-26, 10-26 cells for the DMSO, 500 nM and 20 µM inhibitor conditions respectively). Bars represent the mean and SEM. Wilcoxon Ranked sum test was used to compare the replicate means. **d)** Average size of IRAK1 assemblies. Violin plots show the distribution of individual cell measurements for DMSO, 500 nM and 20 µM kinase inhibitor treatment. The dots superimposed on the violin plots represent the mean IRAK4 assembly size of each replicate (n = 8, 3 and 5 replicates and 5036, 1492 and 2882 assemblies measured all replicates for the DMSO, 500 nM and 20 µM inhibitor concentration respectively). Bars represent the mean and SEM. Wilcoxon Ranked sum test was used to compare the replicate means. **e)** MyD88:IRAK1 complex lifetime (seconds) in cells treated with DMSO (top, N = 910 MyD88:IRAK1 complexes), 500 nM kinase inhibitor (middle, N = 3021 MyD88:IRAK1 complexes) and 20 µM kinase inhibitor (bottom, N = 4387 MyD88:IRAK1 complexes) from a single replicate. **f)** Myddosomes were immunoprecipitated from cell lysates treated without or with IRAK4 kinase inhibitor and analyzed by western blot. Treatment with the IRAK4 kinase inhibitor reduced co-assembly and the post-translational modification of IRAK1 within Myddosomes.

IRAK4 phosphorylation upregulates its kinase activity, and allows IRAK4 to phosphorylate IRAK1 at a threonine residue within its kinase active site^25^. This upregulates IRAK1 kinase activity, leading to extensive IRAK1 autophosphorylation^7^. Additionally, IRAK1 is modified by ubiquitination^26^ in response to IL-1R stimulation and stimulation of the Toll-Like receptors (TLR), which also signal via Myddosomes. IRAK1 incorporation into Myddosome would bring IRAK4/1 into proximity, permitting IRAK4 to phosphorylate and activate further downstream posttranslational modifications of IRAK1. Therefore, one consequence of IRAK4 kinase inhibition would be a lack of phosphorylated or ubiquitinated IRAK1 associated with Myddosomes. To test this and to confirm that IRAK4 kinase inhibition reduces IRAK1 incorporation into Myddosomes, we used co-immunoprecipitation of MyD88-GFP to isolate Myddosomes from IL-1 stimulated cells (Fig. 5F, Supplementary Fig. 4B). Compatible with the reduction in colocalized MyD88:IRAK1 puncta in the inhibitor-treated cells (Fig. 5C), IRAK4 kinase inhibitor treatment reduced IRAK1 association with isolated Myddosomes (Fig. 5F). In inhibitor-treated cells, IRAK1 ran as a single band, suggesting not only less IRAK1 but also unmodified IRAK1. In contrast, in Myddosomes isolated from control cells, IRAK1 ran as a high molecular weight band and smear, consistent with Myddosomes containing hyper-phosphorylated or ubiquitinated IRAK1^7,26^. We conclude that IRAK4 phosphorylation triggers IRAK1 incorporation into the Myddosome and activates its posttranslational processing. Thus, IRAK4 autophosphorylation functions as a regulatory switch that controls downstream signal transduction reactions by regulating DD interactions between IRAK4 and IRAK1.

### IRAK4 autophosphorylation acts as a regulatory switch by potentiating IRAK4/1 death domain co-assembly

Our data suggests that IRAK4 kinase activity and autophosphorylation functions as a regulatory switch that gates heterotypic DD interactions between IRAK4 and IRAK1 and in turn IRAK1 incorporation into Myddosomes. Previously, it has been shown in vitro that the phosphorylated IRAK4 kinase domain forms a hetero-dimer with the IRAK1 kinase domain^27^. To investigate how IRAK4 auto-phosphorylation functions as a switch that controls IRAK4:IRAK1 DD interactions, we directly imaged the co-assembly of IRAK1 with WT and mutant alleles of IRAK4. We reconstituted a MyD88/IRAK4/IRAK1 triple KO EL4 cell line (referred to as 3xKO cells, Supplementary Fig. 5A) with MyD88, IRAK1-mScarlet, WT or mutant alleles of IRAK4 tagged with GFP. We found that both IRAK4^DD^ and IRAK4^K213/214A^ mutants were unable to recruit and incorporate IRAK1 in nascent Myddosome complexes (Fig. 6A, C, Movie S6). While 3.97% IRAK4^WT^ assemblies colocalized stably with IRAK1 (>30s), only 0.28 % and 0.46 % of IRAK4^DD^ and IRAK4^K213/214A^ assemblies stably colocalized with IRAK1 (Fig. 6D). This result is consistent with pIRAK4 kinase domain directly interacting with the IRAK1 kinase domain^27^, thereby recruiting it to the nascent Myddosome assembly (Fig. 6E). We conclude that the interaction between the pIRAK4:IRAK1 kinase domains stimulate IRAK1 incorporation and oligomerization of its DD at the MyD88:IRAK4 complex. Thus autophosphorylation is an essential regulatory switch that activates the final stages of Myddosome assembly (Fig. 2E) and thereby exerts control over downstream inflammatory signaling.

**Figure 6.**
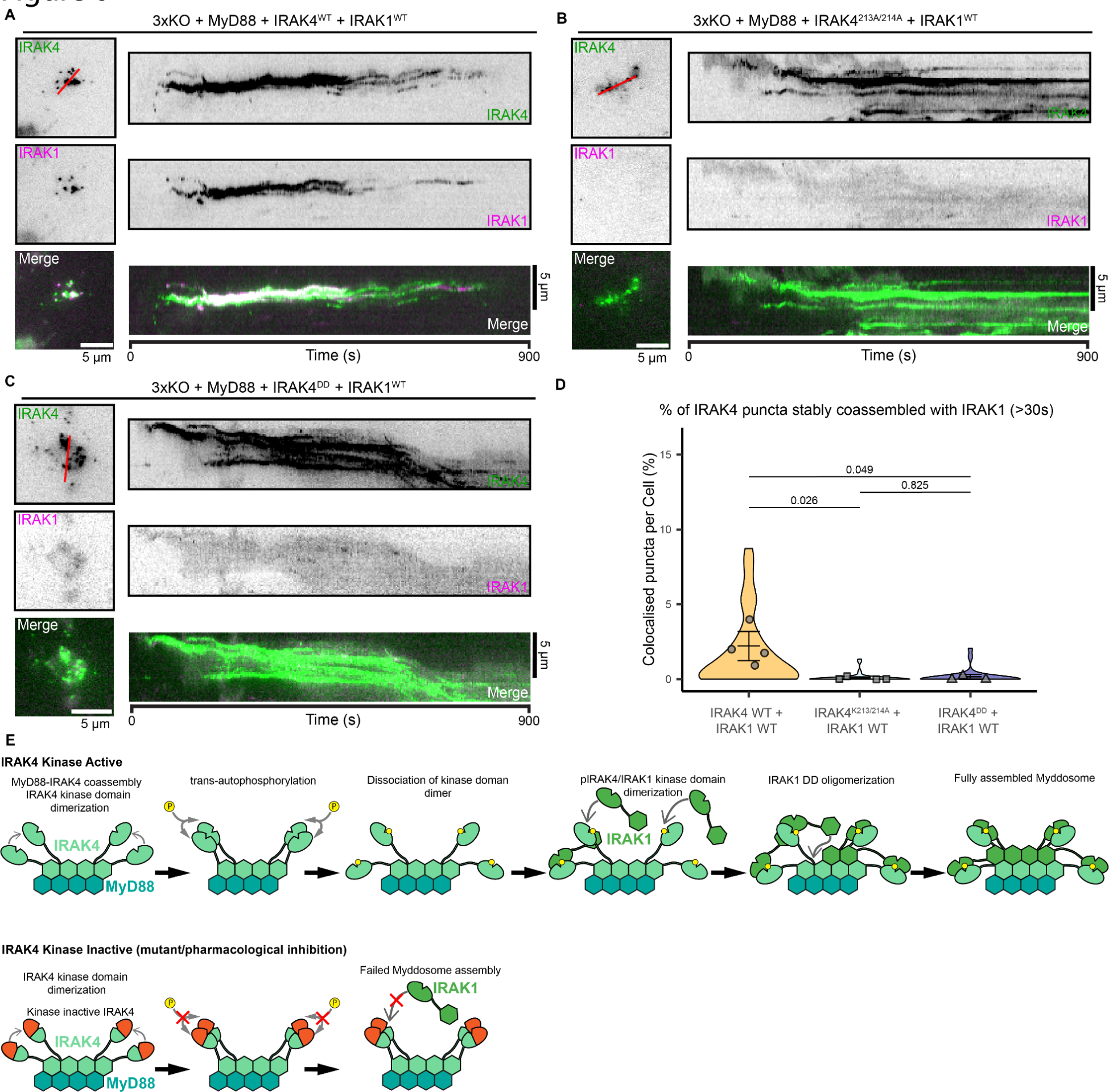
IRAK4 kinase domain autophosphorylation is essential for co-assembly with IRAK1 **a-c)** TIRF images and kymograph analysis of EL4-MyD88/IRAK4/IRAK1-KO (3xKO) reconstituted with MyD88 and a combination of IRAK4^WT^-GFP/IRAK1^WT^-mScarlet (a) or IRAK4^K213A/K214A^-GFP-IRAK1^WT^-mScarlet (b) or IRAK4^DD^ GFP/IRAK1^WT^-mScarlet (c). Kymographs derived from red line overlaid TIRF images. Scale bar, 5 µm. **d)** Percentage of IRAK4 puncta that colocalize with IRAK1 per cell. Violin plots show the distribution of individual cell measurements for 3xKO cells reconstituted with IRAK1^WT^ and either IRAK4^WT^ or IRAK4^K213/214A^or IRAK4^DD^. The dots superimposed on the violin plots represent the mean value of each replicate (n = 4, 4 and 3 replicates with each replicate encompassing measurements from 5-15, 7-18, 4-18 cells for the IRAK4^WT^, IRAK4^K213/214A^ or IRAK4^DD^ expressing cells respectively). Bars represent the mean and SEM. Wilcoxon Ranked sum test was used to compare the replicate means. **e)** Schematic model of how IRAK4 autophosphorylation is a switch that activates IRAK1 death domain assembly and incorporation into Myddosomes. IRAK4 kinase domain autophosphorylation disrupts the homo-dimer of IRAK4 kinase domain, and allows it to hetero-dimerize with the IRAK1 kinase domain. This interaction tethers IRAK1 at assembling Myddosomes thereby facilitating IRAK1 DD oligomerization. This triggers the oligomerization of IRAK1 death domain at nascent Myddosomes.

### IRAK4 autophosphorylation is a universal switch that controls incorporation of IRAK1/2/3 into the myddosome

To determine whether the role of IRAK4 autophosphorylation extends beyond the control of IRAK1 incorporation we looked at IRAK2 and IRAK3. Similar to IRAK1, IRAK2 and IRAK3 are also able to co-assemble with MyD88:IRAK4. IRAK2 is able to activate an equivalent downstream inflammatory signaling reaction as IRAK1^9^, suggesting functional redundancy. In contrast IRAK3 is proposed to be a negative regulator in signal transduction^11^. IRAK2 and IRAK3, unlike IRAK1, are putative pseudokinases^5,6^. Given these biochemical differences, it is possible that the Myddosomes composed of either IRAK1, IRAK2 or IRAK3 could have different mechanisms of assembly. However, if IRAK4 auto-phosphorylation is a universal regulatory switch activating IRAK DD assembly, it would control the co-assembly of IRAK2 or IRAK3 into Myddosomes. We found that EL4 cells do not express detectable levels of IRAK2 or IRAK3 (Fig. S6A). Therefore to examine the formation of IRAK2 and IRAK3 containing Myddosomes, we reconstituted EL4-MyD88-GFP IRAK1KO^-/-^ (IRAK1 KO) cell line^17^ with murine IRAK2-mScarlet or IRAK3-mScarlet . We found that IRAK2 and IRAK3 are incorporated into Myddosome with similar kinetics to IRAK1 (Supplementary Figure 6B-H, Movie S7 and S8).

To test whether IRAK4 autophosphorylation regulates IRAK2/3 incorporation, we imaged both in the presence of DMSO or IRAK4 kinase inhibitor (Fig. 7A-D, Supplementary Fig. 7A, B). We found that treatment with IRAK4 kinase inhibitor decreased the percentage of MyD88 puncta that recruited IRAK2/3, with the most potent effect observed at 20 µM of IRAK4 inhibitor (Fig. 7E). Similar to IRAK1, the IRAK4 kinase inhibitor decreases the size of IRAK2/3 assemblies, and the lifetime of MyD88:IRAK2/3 colocalized puncta (Fig. 7F-G). We conclude that IRAK4 kinase activity and autophosphorylation regulates IRAK1/2/3 incorporation and DD oligomerization at Myddosomes. Therefore, IRAK4 autophosphorylation is a regulatory switch that controls the final assembly stage of Myddosomes, regardless of the final composition.

**Figure 7.**
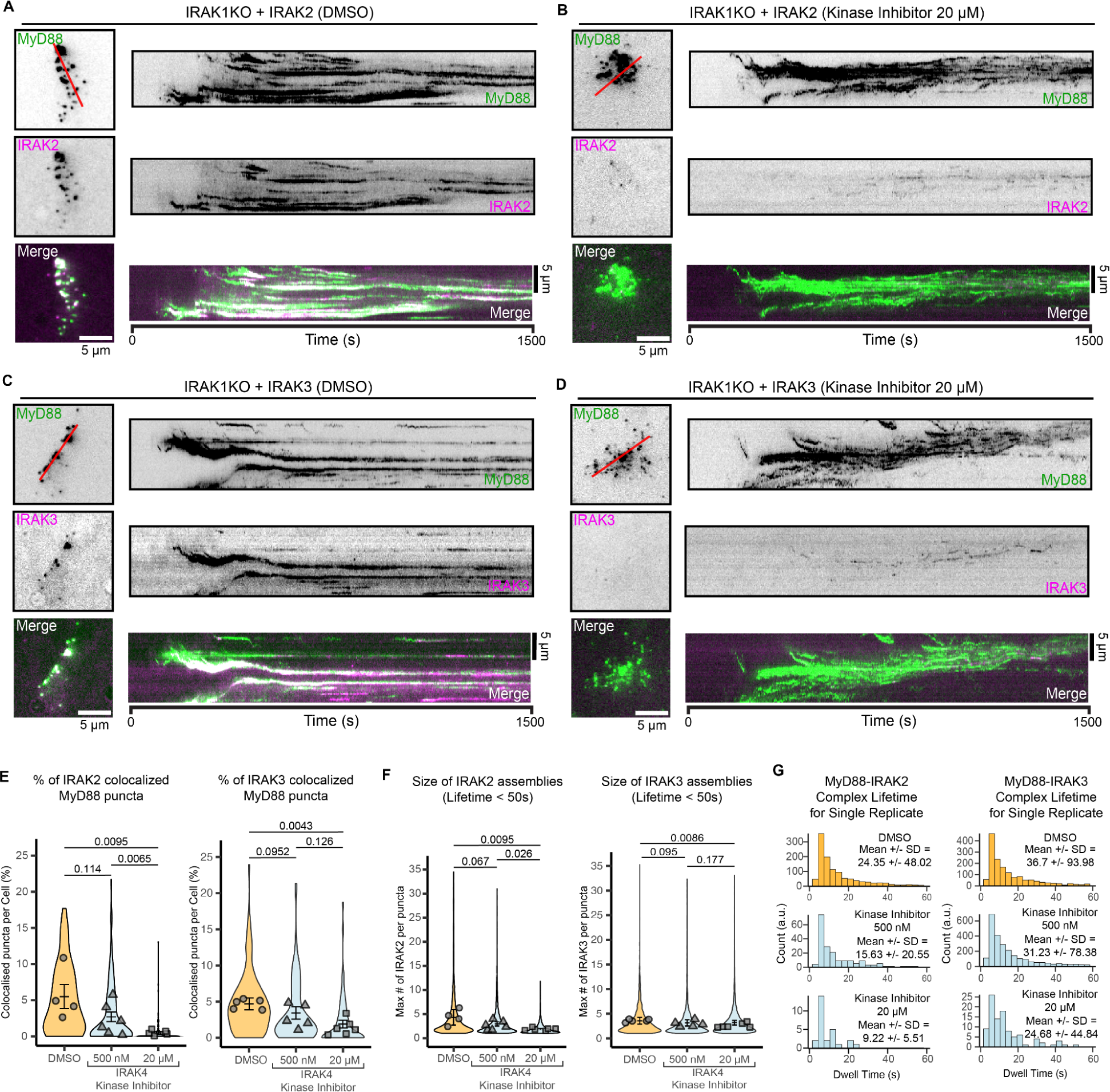
Inhibition of IRAK4 kinase activity disrupts the incorporation of IRAK2/3 into nascent Myddosome complexes. **a-d)** TIRF images and kymograph analysis of DMSO control or IRAK4 kinase inhibitor treated EL4-MyD88-GFP/IRAK1-KO reconstituted with IRAK2-mScarlet (a-b) or IRAK3-mScarlet (c-d). Kymographs derived from red line overlaid TIRF images. Scale bar, 5 µm. **e)** Percentage of MyD88 puncta that colocalize with IRAK2 (left panel) or IRAK3 (right panel) per cell in DMSO control or IRAK4 inhibitors treated cells. Violin plots show the distribution of individual cell measurements for DMSO, 500 nM and 20 µM kinase inhibitor treatment. The dots superimposed on the violin plots represent the mean value of each replicate (for IRAK2, n = 4, 6 and 6 replicates with each replicate encompassing 14-27, 16-33, 15-35 cells for the DMSO, 500 nM and 20 µM inhibitor conditions respectively; for IRAK3 n = 5, 5 and 6 replicates for the DMSO, 500 nM and 20 µM inhibitor conditions respectively, each replicate encompasses measurements from 5-30, 8-29 and 5-35 cells). Bars represent the mean and SEM. Wilcoxon Ranked sum test was used to compare the replicate means. **f)** Average size of IRAK2 (left panel) or IRAK3 (right panel) assemblies in DMSO control or IRAK4 inhibitor treated cells. Violin plots show the distribution of individual cell measurements for DMSO, 500 nM kinase inhibitor treatment and 20 µM kinase inhibitor treatment. The dots superimposed on the violin plots represent the mean IRAK4 assembly size of each replicate (for IRAK2, n = 4, 6, and 6 replicates and 2050, 2890 and 2074 assemblies measured across all replicates for the DMSO, 500 nM and 20 µM inhibitors condition respectively, for IRAK3, n = 5, 5 and 6 replicates and 3419, 3449, and 3288 assemblies measured all replicates for the DMSO, 500 nM and 20 µM inhibitors condition respectively). Bars represent the mean and SEM. Wilcoxon Ranked sum test was used to compare the replicate means. **g)** MyD88:IRAK2 (left panel) and MyD88:IRAK3 (right panel) complex lifetime (seconds) in cells treated with DMSO (top, N = 1217 MyD88:IRAK2 complexes and N = 1710 MyD88:IRAK3 complexes), 500 nM kinase inhibitor (middle, N = 189 MyD88:IRAK2 complexes and 2723 MyD88:IRAK3 complexes), and 20 µM kinase inhibitor (bottom, N = 27 MyD88:IRAK2 complexes and 109 MyD88:IRAK3 complexes), from a single replicate.

## Discussion

Here, we demonstrate that autophosphorylation of the IRAK4 kinase domain plays a central role in IL-1R-activated Myddosome assembly. By reconstructing a timeline of Myddosome assembly, we find that IRAK4 autophosphorylation is a distinct step that occurs after MyD88:IRAK4 assembly but before the incorporation of IRAK1 (Fig. 2A-E, Supplementary Fig. 2, Fig. 3A-E, Fig. 4A-G). This step is critical because it potentiates the recruitment and activation of IRAK1 (Fig. 5). Finally, we show that this mechanism is universal because it also controls IRAK2/3 incorporation (Fig. 7). Thus, IRAK4 autophosphorylation is a biochemical switch that controls the DD assembly of IRAK1/2/3, which is the final stage of Myddosome assembly and required for downstream signaling. Our data shows the assembly of a DD signaling complex is not solely controlled by oligomerization. Instead, DD assembly can integrate a powerful regulatory switch using an energy dependent phosphorylation mechanism to control signal transduction.

A critical question is how IRAK4 autophosphorylation universally regulates IRAK1/2/3 incorporation. From structural studies, unphosphorylated IRAK4 kinase domains form a homodimer, and this interaction is broken by trans-autophosphorylation^13^. Phosphorylated IRAK4 kinase domains interact with the kinase domains of IRAK1/2/3^6,27^. One possibility is that IRAK4 kinase domain phosphorylation functions as a switch by converting the IRAK4 kinase domain homodimer into a heterodimer of IRAK4:IRAK1/2/3 kinase domain (Fig. 6E). This step is critical, because our data shows that IRAK4 kinase domain mutants are incapable of co-assembling with IRAK1 (Fig. 6). Therefore, the heterodimer of the IRAK kinase domains possibly tethers IRAK1/2/3 at nascent Myddosome and stimulates DD oligomerization.

Beyond IL-1R signaling, multiple inflammatory signaling pathways transduce signals through oligomeric complexes composed of DDs^28,29^. Our data shows that multimerization of DDs can be controlled by other biochemical processes, such as post translational modification, beyond the specific interactions between these domains. The integration of autophosphorylation and death domain oligomerization stages is possibly conserved outside of mammals, as Toll receptor activation triggers autophosphorylation of Drosophila IRAK homolog Pelle kinase^30^. Similar to IRAKs, we find enzymatically active signaling proteins that contain death domains and assemble into multimeric complexes, for example receptor-interacting protein kinases (RIPK)^31–33^. This suggests broader prevalence of enzymatic activity controlling protein oligomerization and signaling cascades. Our insight into hierarchical interplay between macromolecular assembly and enzymatic activity explains why IRAK4 kinase inhibitors disrupt controlled signal transduction. Therefore, this points to a possible broader therapeutic strategy of targeting enzymatic activity to disrupt macromolecular assembly and consequently signal transduction.

## Acknowledgements

We thank Vikram Rao and Pfizer for providing a specific antibody against phospho-IRAK4. We thank the Max Planck Computing and Data Facility for providing cluster computer resources for data analysis. We thank Kathrin Laettig for assistance with ELISA and cell culture. We thank all Taylor lab members for critical feedback and commenting on the project. We thank Iain Patten for assistance in drafting and constructing the manuscript.

This work was supported by the Max Planck Society and the DFG (project number 499533619). The authors declare no competing financial interests and no conflicts of interest.

## Author contributions

M.J. Taylor, N. Srikanth and E. Ziska, purified proteins for labeling membranes. N. Srikanth and R. Deliz-Aguirre wrote and validated custom image analysis scripts. N. Srikanth performed imaging experiments. E. Ziska performed Western Blot verification of cell lines and time course analysis of pIRAK4 production. E. Ziska and D. Gola performed immunoprecipitation and Western Blotting experiments. N. Srikanth and M.J Taylor performed experimental design. N. Srikanth performed data analysis and visualization. M. Bilay performed data analysis on blinded pIRAK4 immunofluorescence data. The project was conceived and supervised by M.J. Taylor. The manuscript was prepared by N. Srikanth, E. Ziska and M.J. Taylor with input from all authors.

## Material and Methods

### KEY RESOURCES TABLE

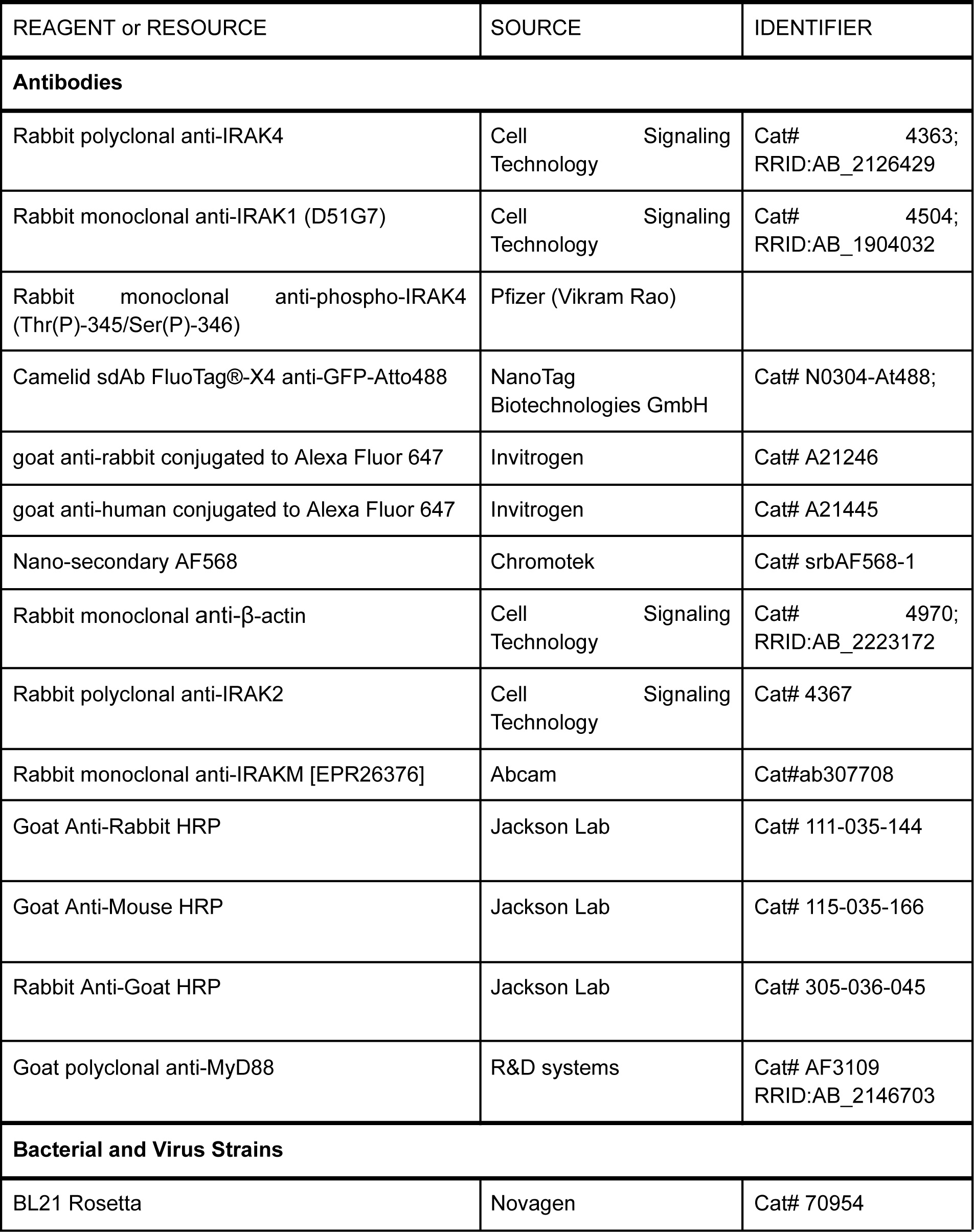
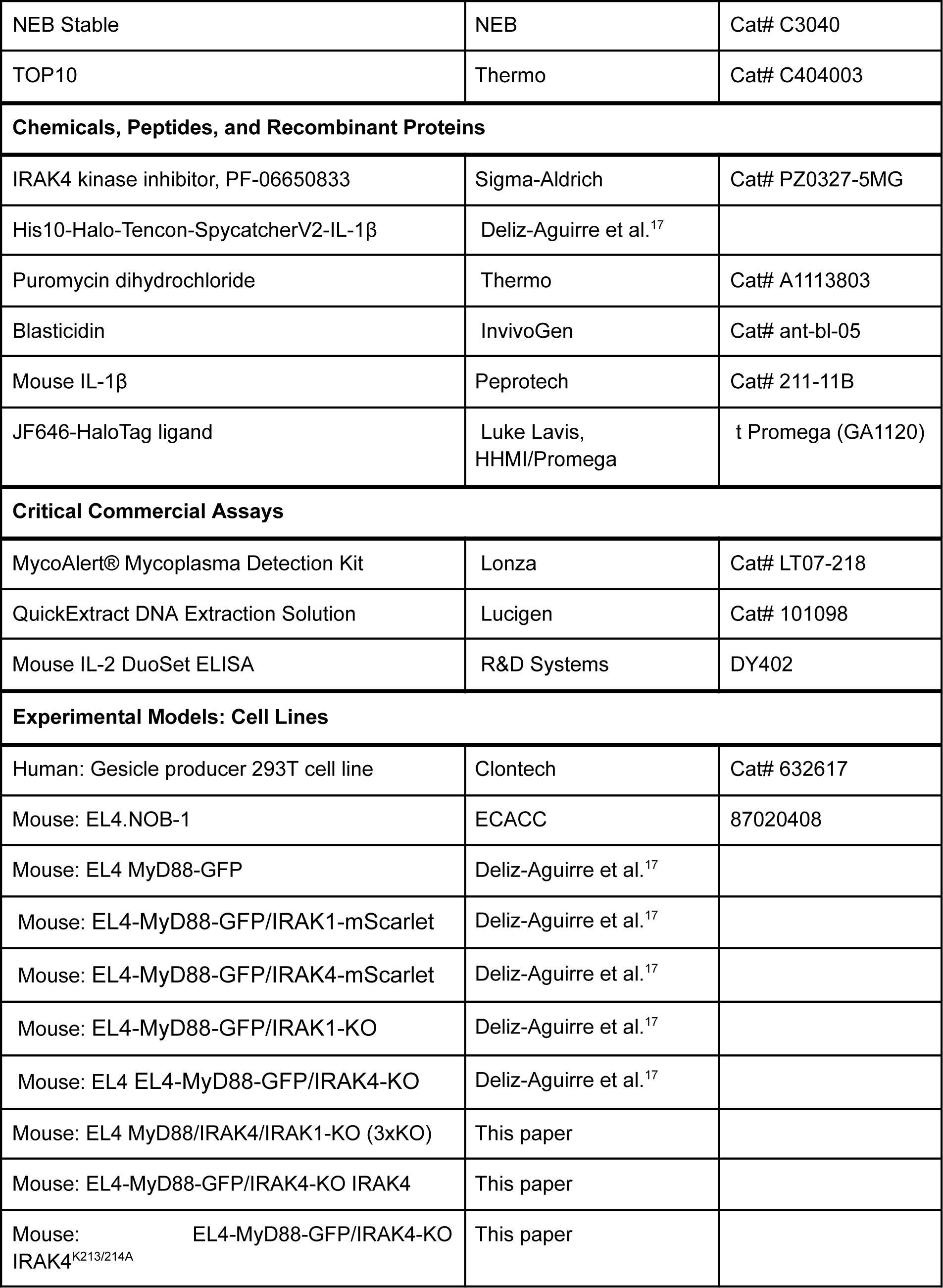
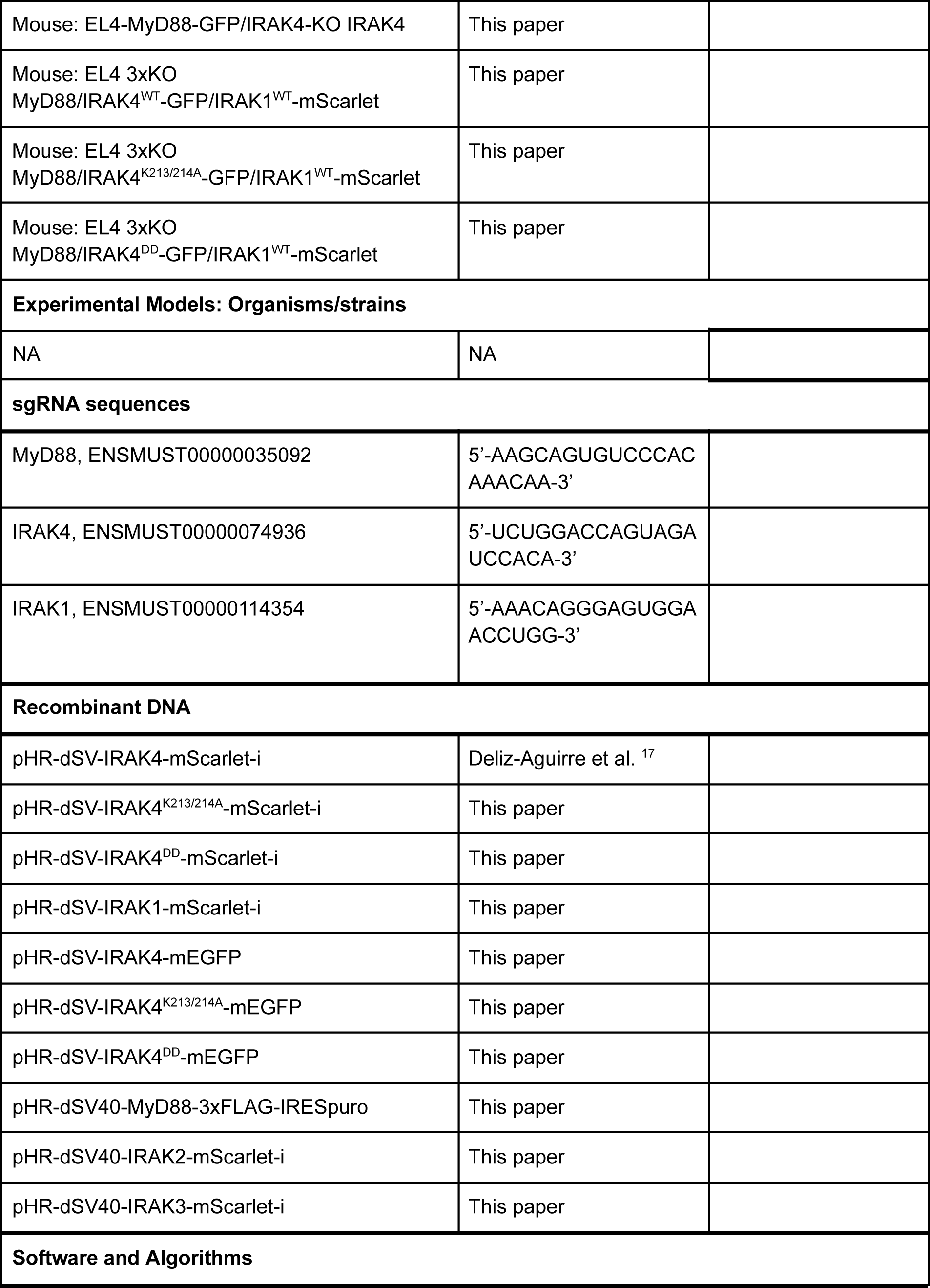
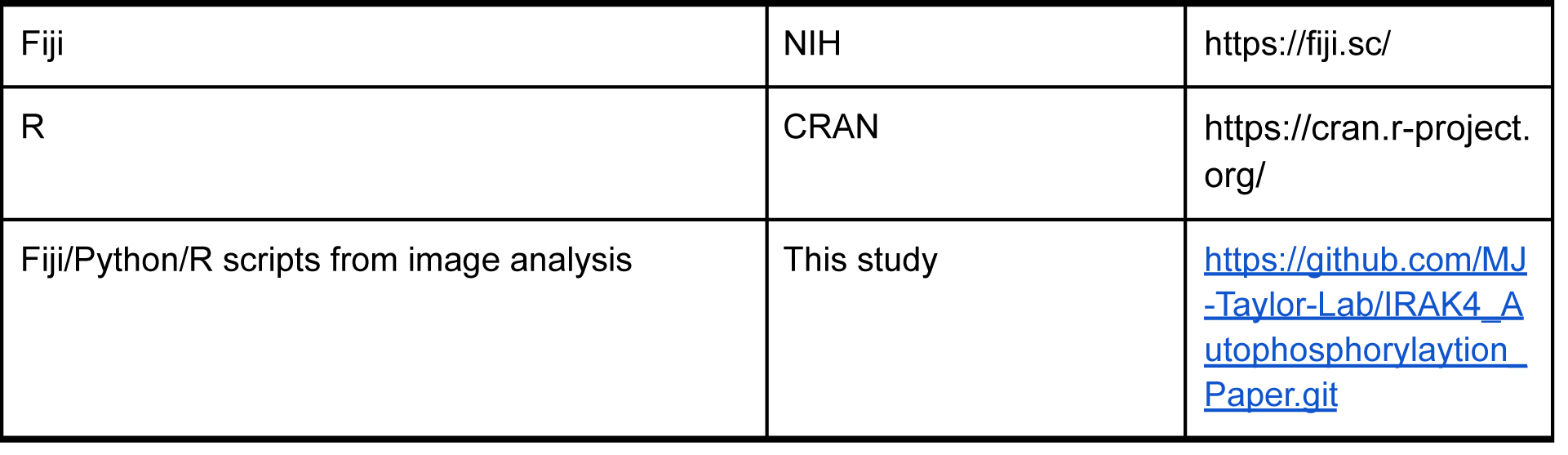

## Methods

### Cell Culture

EL4.NOB1 WT (ECACC, and referred to as EL4 in the paper) and gene-edited cells were grown in RPMI (Thermo Fisher Scientific) with 10% FBS (Sigma) supplemented with 2 mM L-glutamine. EL4 cultures were maintained at a cell density of 0.1-0.5x 10^6^ cells/ml in 5% CO2, 37°C. HEK-293T cells (Clontech) were grown in DMEM (Thermo Fisher Scientific) with 10% FBS supplemented and 2 mM L-glutamine. All cells were determined to be negative for mycoplasma using the MycoAlert detection kit (Lonza).

### Generation of CRISPR/Cas9 triple MyD88/IRAK1/IRAK4 knockout Cell Lines

We purchased WT EL4.NOB1 cells that had been electroporated with Cas9 RNPs loaded with sgRNA targeting the MyD88 (gRNA sequence 5’-AAGCAGUGUCCCACAAACAA-3’, ENSMUST00000035092) IRAK4 (gRNA sequence 5’-UCUGGACCAGUAGAUCCACA-3’, ENSMUST00000074936) and IRAK1 (gRNA sequence 5’-AAACAGGGAGUGGAACCUGG,

ENSMUST00000114354) gene locus from the Synthego Corporation (Redwood CA, USA). From this pooled population, we isolated single cell clones by limited dilution on 96 well plates. We screened and identified MyD88/IRAK1/IRAK4 triple knockout single cell clones by western blot analysis (Supplementary Fig. 5A). Western blot analysis was performed with the following antibodies: rabbit polyclonal anti-IRAK4 (Cell Signaling Technology, #4363), rabbit monoclonal anti-IRAK1 (D51G7, Cell Signaling Technology, #4504), goat polyclonal anti-MyD88 (R&D systems, #AF3109), and Mouse monoclonal anti-β-actin (Cell Signaling Technology, #4970). EL4 MyD88/IRAK4/IRAK1-KO knockout cell lines referred to as EL4-3xKO.

### Western blot analysis of IRAK4 phosphorylation

EL4 wt cells were seeded into 6 well plates at a density of 7.5x10^5^ cells/well 24 hours before stimulation. On the next day cells were stimulated with 1ng/ml of recombinant mouse IL-1β (Peprotech) in RPMI for several time points between 1-120 min. To prepare cell lysates, the stimulated and unstimulated cells were briefly washed in PBS to remove IL-1, and then resuspended in 2x SDS-PAGE lysis buffer (Thermo 15493939). Lysates were incubated for 5 mins at 95 C before run on NuPAGE Novex 4-12% Bis-Tris Protein Gels (Thermo) . Samples were then analyzed by western blotting. To visualize the pIRAK signal intensity relative to the IRAK4 signal, the blots were developed with Rabbit monoclonal anti-pIRAK4 (Thr(P)-345/Ser(P)-346) (a gift from Vikram Rao, Pfizer ^21^), Rabbit polyclonal anti-IRAK4 (Cell Signaling Technology, Cat# 4363) and Mouse monoclonal anti-ß-actin (Cell Signaling Technology, #4970) for normalization. Chemiluminescence detection (Super signal substrate, Pierce #34095) was imaged with the BioRad ChemiDoc MP Imaging System. For each replicate we ran two gels and WB blots loaded with the identical amount of sample, for each replicate one gel was blotted for IRAK4 and the other for pIRAK4 (see Supplementary Fig. 1A). After IRAK4 or pIRAK4 detection, both blots were stripped and then reprobed with anti-actin to normalize the signal intensity of respective IRAK4/pIRAK4 signal between the two gels. We performed analysis of the Western blot band intensity and quantified the production of pIRAK4 in ImageJ/Fiji. The intensity of the IRAK4 and pIRAK4 band was normalized to the actin loading control on each blot for each replicate. The normalized pIRAK4 value was then divided by the normal IRAK4 value to give a relative value of pIRAK4 production at each time point assayed. Data was visualized and plotted in R.

For Western Blot analysis of pIRAK4 levels in IRAK4 kinase inhibitor treated/untreated EL4 cells (Supplementary Fig. 2B), we incubated EL4 cells with media containing 20 µM of IRAK4 kinase inhibitor (PF-06650833, stock concentration of 10 mM in DMSO) for 4 hours. DMSO at a final concentration of 0.2% v/v was added to the control samples. EL4 cells were then stimulated with 10-100 ng/ml of recombinant mouse IL-1β for 30 mins. We prepared cell lysates as described above, and ran them on NuPAGE Novex 4-12% Bis-Tris Protein Gels (Thermo). After membrane transfer, cell lysates were probed with anti-pIRAK4 and developed by chemiluminescence as described above.

### Co-Immunoprecipitation of Myddosomes from EL4 Cell Lysates

24 hours before stimulation, 5x10^6^ EL4 MyD88-GFP cells were seeded in T-75. The next day, the medium was removed and fresh medium, containing 20µM IRAK4 kinase inhibitor or 0.2% DMSO only, was added. The cells were incubated with IRAK4 kinase inhibitor or DMSO at 37°C for 4 hours. The cells were stimulated with 100 ng/ml of recombinant mouse IL-1β for 15 mins. Stimulated and unstimulated cells were then washed in ice cold PBS, spun down and resuspended in lysis buffer (50 mM Tris, 150 mM NaCl, 1% Igepal/ NP40, pH 8.0). Lysis buffer contained phosphatase and protease inhbitiors (PhosSTOP, Roche, Cat# 04906837001 and Protease Inhibitor Mini Tablets, EDTA-free, Pierce, Cat# A32955) at the manufacturer’s recommended concentration. Aliquots of the lysates were stored for downstream analysis (input fraction). Lysates were added to GFP beads (Chromotek: GFP-Trap Magnetic Particles, Cat# M-270) pre-equilibrated in Lysis buffer, and incubated overnight with rotation at 4°C. The next day beads were applied to magnetic racks and the supernatant was removed. Next, the beads were washed in Lysis buffer, and eluted in 50 µl SDS sample loading buffer. The beads were vortexed and incubated for 5 mins at 95°C. The eluted and input fractions were then ran on NuPAGE Novex 4-12% Bis-Tris Protein Gels (Thermo) and analyzed by Western Blot using the antibodies and protocols described above (Fig. 5F, Supplementary Fig. 4B)

### Generation of lentiviral constructs

Full details of the Lentiviral plasmid construction are given below. All full length constructs were confirmed by Sanger sequencing.

#### pHR-dSV-IRAK4^K2^^13/14A^-mScarlet-i/mEGFP, pHR-dSV-IRAK4^DD^-mScarlet-i/mEGFP

To prepare the pHR-dSV-IRAK4^K213/214A^-mScarlet-i and pHR-dSV-IRAK4^DD^-mScarlet-i constructs, the mouse IRAK4 sequence with two mutations (K213/214A) and the mouse IRAK4 sequence excluding the kinase domain were ordered as gBlocks (IDT). These gBlocks were fused with a PCR fragment of mScarlet-i (amplified by PCR using the forward

5’-tgctacaagagatgtctgctggaggaagtggaggttctggtggtagtgtg-3’ and reverse 5’-aggtcgactctagagtcgcggcc-3’ primers) or mEGFP (PCR using the forward 5’-tgctacaagagatgtctgctggaggaagtggaggttctggtggtagtgtg-3’ and reverse 5’-ggccgcgactctagagtcgacct-3’ primers) and Mlu1/Not1 digested pHR-dSV lentiviral plasmid using Gibson assembly 34. IRAK4 constructs were fused in frame to mScarlet-i or mEGFP via a 3xGGS linker.

#### pHR-dSV-IRAK4-mEGFP

Full length IRAK4 (BC051676) was amplified by PCR from a previously described ^17^ pHR-dSV-IRAK4-mScarlet-i construct. mEGFP was amplified using the above-mentioned primers. DNA fragments were assembled with pHR-dSV lentiviral plasmid digested with Mlu1/Not1 using Gibson Assembly.

#### pHR-dSV-IRAK1-mScarlet-i

Full length Mouse IRAK1 (NCBI Reference Sequence: NP_001171445.1) was ordered as a codon optimized gBlock (IDT). The gBlock encoded a complementary 5’ end to pHR-dSV. This DNA fragments was fused to PCR fragment of mScarlet-i (amplified by PCR using the forward 5’-gggggtagcggaggttctggtggtagtgtgagcaaggg-3’ and reverse 5’-cctgcaggtcgactctagagtcgcggccgctttacttgtacagctcgtccatgcc-3’ primers) and a pHR-dSV lentiviral plasmid digested with Mlu1/Not1 using Gibson Assembly. IRAK1 was fused in frame to mScarlet-i via a 3xGGS linker

#### pHR-dSV-IRAK2-mScarlet-i, pHR-dSV-IRAK3-mScarlet-i

Full length Mouse IRAK2 (NCBI Reference Sequence: NP_751893.3) and full length Mouse IRAK3 (NCBI Reference Sequence: NC_000076.7) was ordered as a codon optimized gBlock (IDT). The gBlock encoded a complementary 5’ end to pHR-dSV. This DNA fragment was fused to the pHR-dSV-IRAK1-mScarlet-i lentiviral plasmid digested with Mlu1/BamH1 (to excise the IRAK1 fragment) using Gibson Assembly. IRAK2/3 was fused in frame to mScarlet-i via a 3xGGS linker.

#### pHR-dSV40-MyD88-3xFLAG-IRES-Puro

Mouse MyD88 fused to 3xFLAG-IRES-Puro was ordered as a codon optimized gBlock (IDT). The gBlock encoded a complementary 5’ end to pHR-dSV. This DNA fragment was fused to the pHR-dSV-IRAK1-mScarlet-i lentiviral plasmid digested with Mlu1/Not1 (to excise the IRAK1 mScarlet-i fragment) using Gibson Assembly.

### Lentiviral Production and Generation of Stable Expressing EL4 Cell Lines

Lentivirus particles were produced in HEK-293T cells by co-transfection of the pHR transfer plasmids with second-generation packaging plasmids pMD2.G and psPAX2 (a gift from Didier Trono, Addgene plasmid # 12259 and # 12260). Virus particles were harvested from the supernatant after 48-72 hrs, filtered, and applied to EL4 cells. After three days, the transduced EL4 cells were resuspended in fresh RPMI medium. After lentiviral transduction, we select positive EL4 cells expressing fluorescent protein fusion of IRAK4/1 using a BD FACS Aria II at the Deutsches Rheuma-Forschungszentrum Berlin Flow Cytometry Core Facility. Specific details of EL4 cell line generation given below.

To engineer EL4 cells expressing IRAK4 alleles, we transduced EL4-MyD88-GFP/IRAK4 KO cells with lentiviruses encoding WT or mutant IRAK4. 1-2 weeks after lentivirus transduction, we used FACS to select cell populations with homogenous expression levels. EL4 WT cells were used as a negative control and the EL4-MyD88-GFP/IRAK4-mScarlet knock-in cell line was used to create a sorting gate that selected transduced cells with expression levels that were equivalent to endogenously expressed IRAK4.

To engineer EL4 cells expressing IRAK2 or IRAK3 tagged with mScarlet, we transduced EL4-MyD88-GFP/IRAK1 KO cells with lentiviruses encoding WT IRAK2 or IRAK3 fused to mScarlet-i. 1-2 weeks after lentivirus transduction, we used FACS to select cell populations with homogenous expression levels. EL4 WT cells were used as a negative control and the EL4-MyD88-GFP/IRAK1-mScarlet knock-in cell line was used to create a sorting gate that selected transduced cells with expression levels that were equivalent to endogenously expressed IRAK1.

To engineer the EL4-3xKOs EL4 expressing MyD88, IRAK4-GFP and IRAK1-mScarlet, we transduced EL4-3xKOs with lentivirus encoding MyD88-FLAG-IRES-Puromycin. After 1 week of selection with media containing 1 µg/ml of puromycin for 1 week, single cells were sorted in 96 well plates. Single cell clones that grew were assayed by western blot for MyD88 expression level that matched the MyD88-GFP/IRAK4-mScarlet EL4 cell line. Based on this analysis we selected a single clone and transduced it with lentivirus encoding IRAK1^WT^-mScarlet-i and either IRAK4^WT^-mEGFP or IRAK4^K213/14A^-mEGFP or IRAK4^DD^-mEGFP. Cells were then FACS sorted to select GFP/mScarlet positive cells after 1-2 weeks post lentiviral transduction.

### Assay of IL-2 release in WT and gene-edited EL4 cells

To measure IL-2 release, we used the Mouse IL-2 DuoSet ELISA kit (R&D Systems, DY402-05) using the manufacturer’s protocol. The day before IL-1 stimulation, EL4 cells were resuspended into fresh media. In experiments that measured the effect of the IRAK4 kinase inhibitor, this resuspension media contained the inhibitor (or equivalent 0.2% v/v of DMSO-only in untreated samples) at the reported concentration (Supplementary Fig. 2A). The next day, the cells were counted and resuspended in fresh media. We then seeded cells in 48-well plates with 1x10^6^ cells in 150 μl of media per well. Cells were allowed to settle for 30 min, before being stimulated with recombinant mouse IL-1β (PeproTech) in 50 μl medium per well at a final concentration of 5 ng/ mL. For IL-2 ELISA measurements performed with IRAK4 kinase inhibitor treated cells, the concentration of the inhibitor in the media was kept constant throughout the entire duration of the assay. For the unstimulated controls, 50 μl medium was added with/ without the inhibitor. After incubating the cells for 24 h with IL-1, the plates were centrifuged (800x g for 5 minutes), and supernatants were transferred into a new plate. Supernatants were stored at -80°C until IL-2-ELISA analysis. Absorbance readings were acquired on a VersaMax Microplate Reader (Molecular Devices) at 450 nm. IL-2 release was assayed on three independent days in triplicates. The obtained results were expressed as the fold change in IL-2 release between the stimulated and unstimulated samples (Supplementary Fig. 2A, Supplementary Fig. 3A).

### Imaging Chambers and Supported Lipid Bilayers

SLBs were prepared using a previously published method ^19^. Phospholipid mixtures consisting of 97.5% mol 1-palmitoyl-2-oleoyl-sn-glycero-3-phosphocholine (POPC), 2% mol 1,2-dioleoyl-sn-glycero-3-[(N-(5-amino-1-carboxypentyl)iminodiacetic acid)succinyl] (ammonium salt) (DGS-NTA) and 0.5% mol 1,2-dioleoyl-sn-glycero-3- phosphoethanolamine-N-[methoxy(polyethylene glycol)-5000] (PE-PEG5000) were mixed in glass round-bottom flasks and dried down with a rotary evaporator. All lipids used were purchased from Avanti Polar Lipids. Dried lipids were placed under vacuum for 2 hrs to remove trace chloroform and resuspended in PBS. Small unilamellar vesicles were produced by several freeze-thaw cycles in conjunction with sonication. Once the suspension had cleared, the lipids were spun in a benchtop ultracentrifuge at 35,000xg for 45 min and kept at 4°C for up to five days.

Supported lipid bilayers (SLB) were formed in 96-well glass bottom plates (SWISSCI). 96-well plates were cleaned for 30 min with a 5% Hellmanex solution containing 10% isopropanol heated to 50°C, then incubated with 5% Hellmanex solution for 1 hour at 50°C, followed by extensive washing with pure water. 96-well plates were dried with gas nitrogen and sealed using clear Polyolefin sealing tape (Thermo Scientific) until needed. To prepare SLB on 96-well plates, individual wells were cut out and base etched for 15 min with 5 M KOH and then washed with water and finally PBS.

To form SLBs, SUVs suspension was deposited in each well and allowed to form for 1 hr at 45°C. After 1 hr, wells were washed extensively with PBS. SLBs were incubated for 20 min with HEPES buffered saline (HBS: 20 mM HEPES, 135 mM NaCl, 4 mM KCl, 10 mM glucose, 1 mM CaCl2, 0.5 mM MgCl2) containing with 5 mM NiCl2 to charge the DGS-NTA lipid with nickel. The SLBs were then washed in HBS containing 0.1% BSA to block the surface and minimize non-specific protein adsorption and incubated at room temperature for 30 min. For SLBs set up on 96-well plates the total well volume was 690 μl. Each well was completely filled with HBS containing 0.1% BSA, and 570 μl was removed leaving 120 μl in each well. After blocking, the SLBs were functionalized by incubation for 1 hr with 120 µl His-tagged proteins to get a final concentration of 5 nM. The labeling solution was then washed out with HBS.

### Protein Expression, Purification and Labeling

We functionalized supported membranes with an engineered variant of mouse IL1β (in text referred to as His10-Halo-JF646-IL-1β). This IL-1 variant had been engineered to have a 10x His tag, and could therefore label membranes that contained DGS-NTA lipids. Full details on the design, expression and purification of this construct have been previously reported (Deliz et al., 2021 and Cao et al., 2023). For microscopy calibration of mScarlet single molecule intensity we used His10-mScarlet-IL-1β (previously described here: Deliz-Aguirre et al., 2021). For mEGFP single molecule intensity calibration, mEGFP was expressed and purified as previously described (see ^17^ and ^35^).

### TIRF-Microscopy data acquisition

Dual color imaging of EL4 cells was performed on an inverted microscope (Nikon TiE, Tokyo, Japan) equipped with NIKON fiber launch TIRF illuminator. Illumination was controlled with a laser combiner using the 488, 561 and 640 nm laser lines. Fluorescence emission was collected through filters for GFP (525 ± 25 nm), RFP (595 ± 25 nm) and JF646 (700 ± 75 nm). All images were collected using a Nikon Plan Apo 100x 1.4 NA oil-immersion objective that projected onto a Photometrics 95B Prime sCMOS camera with 2x2 binning and a 1.5x magnifying lens (calculated pixel size of 0.147 µm). Image acquisition was performed using NIS-elements software. All experiments were performed at 37°C. The microscope stage temperature was maintained using an OKO Labs heated microscope enclosure. Imaging frames of live cells was acquired at an interval of 3 s using exposure times of 200-400 ms.

### Imaging EL4 cells endogenously expressing MyD88-GFP, IRAK4-mScarlet or IRAK1-mScarlet on IL-1β functionalized SLBs with TIRF-Microscopy

His10-Halo-JF646-IL1β functionalized SLBs were set up as described above and as previously described ^17,35^. To quantify the IL-1β density on the SLB before each experiment, wells were prepared that were functionalized with identical labeling protein concentration and time, but with different molar ratios of labelled to unlabelled His10-Halo-IL1β. Before application of cells, SLBs were analysed by TIRF microscopy to check formation, mobility and uniformity. Short time series were collected at wells containing a ratio of labeled to unlabelled His10-Halo-IL1β, (e.g. <1 His10-Halo-JF646-IL1β molecule per µm2) to calculate ligand densities on the SLB based upon direct single molecule counting. All experiments were performed at IL-1 SLB densities of 50 to 80 molecules per µm2.

Before each imaging experiment we acquired calibration images using recombinant mEGFP and mScarlet-i. To image single GFP/mScarlet-i fluorophores, recombinant purified mEGFP and mScarlet-i ^35^ was diluted in HBS and adsorbed to KOH cleaned glass. Single molecules of GFP or mScarlet-i were imaged using identical microscope acquisition settings to those used for cellular imaging. To image live cells, EL4 cells were pipetted onto supported lipids bilayers functionalized with His10-Halo-JF646-IL1β. EL4 cells expressing MyD88-GFP, IRAK4-mScarlet or IRAK1-mScarlet were sequentially illuminated for 70-100 ms with 488 nm and 100ms with 561 nm laser line at a frame interval of 3 s.

For pharmacological inhibition imaging experiments EL4 cells were treated with 500 nM or 20 µM of IRAK4 kinase inhibitor (PF06650833, Sigma) for at least 3 hours before imaging. All treatment conditions contained 0.1% DMSO by volume/volume. Then cells were spun down and resuspended in HBS containing the same concentration of the inhibitor. Inhibitor-treated cells were then added to wells containing HBS with identical concentration of the IRAK4 inhibitor, thereby maintaining the inhibitor concentration through the microscopy acquisition.

### Immunofluorescence staining of phospho-IRAK4

To analyze the colocalization of phospho-IRAK4 with MyD88-GFP and IRAK4-mScarlet or IRAK1-mScarlet (Fig. 2), EL4 cells were stimulated with IL-1β-functionalized SLBs for 30 mins, and then fixed with 4% (wt/vol) PFA containing 0.5% (wt/vol) Triton X-100 for 20 min at room temperature. Staining was then performed with a traditional two-step staining method. After fixation, cells were washed with PBS, then blocked in PBS containing 10% (wt/vol) BSA and 0.1% Triton X-100 at 4° C overnight. The next day, the cells are stained using a 1 step labeling procedure with antibody solution containing anti-pIRAK4 (1:400, Vikram Rao, Pfizer), Nano-Secondary 647 (1:1000, ChromoTek alpaca anti-human IgG/anti-rabbit IgG, recombinant VHH, Alexa Fluor 647 CTK0101, CTK0102 #srbAF647-1) and FluoTag-X4 anti-GFP conjugated to Atto488 (1:500, Nano Tag Biotechnology, #N0304-At488-L) and FluoTag-X4 anti-RFP conjugated to Atto568 (1:500, Nano Tag Biotechnology, #N0404-AF568-L) in PBS containing 10% (wt/vol) BSA and 0.1% Triton X-100 for 1 hr at room temperature. Subsequently, cells were washed with 2 ml 10% (wt/vol) BSA and 0.1% Triton X-100 and finally with 4 ml PBS before imaging with TIRF microscopy.

### Quantification and Statistical Analysis

All data are expressed as the mean ± the standard deviation (SD) or mean ± the standard error of the mean (SEM), as stated in the figure legends and results. The exact value of n and what n represents (such as the number of experimental replicates, cells or MyD88-puncta) is stated in figure legends and results. Means were compared using Wilcoxon Rank-Sum test, due to the small n size of replicate making it difficult to determine whether the data is normally distributed. Data analysis scripts used in this study available at <u>GitHub</u> .

### Quantification of phospho-IRAK4 immunofluorescence

To score MyD88 puncta negative or positive for phospho-IRAK4 and IRAK4-mScarlet or IRAK1-mScarlet, TIRF microscopy images were manually segmented into individual cells. The phosphoIRAK4 and IRAK4-mScarlet or IRAK1-mScarlet channels were blinded, and then MyD88 puncta manually identified and scored negative or positive for colocalization with puncta in the blinded phospho-IRAK4 channel and IRAK4-mScarlet or IRAK1-mScarlet channel. Manual identification and scoring was performed in FIJI with the multi-point annotation tool. We blinded the IRAK4/1-mScarlet and pIRAK4 staining channels before manual scoring of MyD88-GFP puncta. We performed this manual analysis for over 400 MyD88-GFP puncta identified in both EL4-MyD88-GFP/IRAK4-mScarlet and EL4-MyD88-GFP/IRAK1-mScarlet cells. Unblinded manual scoring of MyD88-GFP puncta was visualized as a bar graph (Fig.2b and d).

### Quantification and analysis of Myddosome intensity, lifetime and dynamics

To quantify the dynamics of Myddosome assemblies labeled with MyD88-GFP and IRAK4/1-mScarlet in live EL4 cells, we created an image analysis pipeline that runs in FIJI, Python and R. Image analysis scripts were parallelized so that they may run on cluster computers at the Max Planck Computing and Data Facility (Garching, Bavaria, Germany). The aim of this image analysis pipeline was to first process the images to remove intensity count that derived from camera noise or background fluorescence (non-specific cytosolic fluorescent signal), identify and segment single cells, and then identify and track the individual fluorescently labeled Myddosome complexes within the live cell data. To process the image stack, we first subtracted a dark frame image (i.e., an image only containing intensity values from current and noise generated by the camera electronics) from each image file. The dark frame image was created by calculating the mean intensity from 5000 images acquired without light exposure to the camera, and with identical acquisition settings to those used for live cell imaging.

From this point, dark frame subtracted images were processed to create two sets of image stacks: one set was used for intensity measurements to quantify the dynamics of Myddosome assembly (referred to as the intensity reference image). The second set was processed using different filters and parameters best suited to detect punctate structures, and this was used to optimally identify and track the Myddosomes (referred to as the tracking images). To generate an images stack for tracking Myddosome puncta, the GFP/mScarlet images were processed with a median filter with a 11-pixel diameter. The resulting median filtered image is subtracted from the original to remove background intensity. We next applied two methods to address stochastic fluctuation associated with the fluorophore blinking or camera read noise, both which can contribute to errors in particle detection and linking detected particles between frames. First, we applied a second median blur of 5-pixel diameter to even out the field. Because median filters are edge-preserving (keeps silhouette), a 5-pixel diameter median blur preserved punctate structures while further removing background noise. We then applied a moving average of 3 frames to further suppress intensity fluctuations. Because we were interested in detecting Myddosomes (i.e., puncta that possibly contain both MyD88 and IRAK4/1), we reasoned that combining the GFP and mScarlet channels into a single image would further increase the signal to noise ratio, facilitating the detection and tracking of Myddosome fluorescent puncta. Therefore, all GFP and mScarlet image frames were summed. The result was a single time series with high signal to noise and optimized for particle detection and tracking.

The combination of two median blur filtering steps and running average used to create the tracking reference image skewed intensity values. Therefore, to be able to accurately quantify Myddosome intensity and assembly kinetics, we generated an intensity reference image that preserved intensity values. To accomplish this, we estimate cytosolic background using a median blur with a 25-pixel diameter (3.67µm), which is roughly half the diameter of the cell: SLB interfaces observed in the TIRF field. This median blurred image is an estimate of background intensity from the cytoplasmic fluorescence (that is MyD88-GFP/IRAK4/1-mScarlet not assembled into Myddosome complexes). This estimated background fluorescent intensity is subtracted from the dark frame removed images and the resulting image is the intensity reference image from which all shown and analyzed intensity values are derived.

Next, individual cells within the time series were identified using a marker-controlled watershed segmentation (algorithm from Open Source Computer Vision, OpenCV) implemented on a maximum projection image of the MyD88-GFP intensity channel. The segmented cell boundaries were used to break both the tracking image and intensity reference series into small image series of individual cells. We used the FIJI TrackMate plugin to track the MyD88-GFP/IRAK4/1-Scarlet puncta within the tracking image for each segmented cell. We tracked MyD88-GFP/IRAK4/1-mScarlet puncta with a maximum linkage distance of 5 pixels, a tracking threshold of 1.5 and a tolerance of 2 gap-closing frames between detected particles. After processing in TrackMate, tracking coordinates generated were imported into R. To fill in missing frames (i.e., gap-closing frames with no detected particles) within tracks, we approximated the location of the missing puncta using the Euclidean distance from the coordinates of the preceding and following detected particle. We extracted the fluorescence intensity of puncta from the intensity reference frames of both the MyD88-GFP image and the IRAK4/1-mScarlet image. The intensity was measured from a 5x5 circular pixel region center of the puncta centroid. To achieve sub-pixel accuracy, we segmented a square matrix of pixels from images. This square was centered on the puncta centroid. To reduce computational time, this segmented area was expanded 11 fold with the original pixel intensity subdividing in the expanded area pixels. The square was then multiplied against a binary circle mask (diameter of 5 pixels that was also expanded by 11 fold). The pixel intensities of the product were summed to give a puncta intensity for both the GFP or mScarlet channel.

To estimate the number of MyD88 and IRAK4/1 within tracked Myddosome assemblies, the fluorescent intensity was divided by the median intensity of single GFP and mScarlet fluorophores. To compute the distribution of single fluorophore intensities, images of single GFP or mScarlet fluorophores adsorbed to glass were processed and analyzed identically to MyD88-GFP and IRAK4/1-mScarlet images. We restricted the analysis to MyD88-GFP and IRAK4/1-mScarlet puncta tracked for three or more frames, to focus our analysis to *bona fide* Myddosome assemblies nucleating at the plasma membrane. Puncta were scored positive for IRAK4/1 recruitment when the normalized fluorescent intensity of at least 0.75x IRAK4/1s for 30 seconds (Fig. 3B, Fig. 4D, Fig. 5C and Fig. 7E). IRAK4/1 dwell time distribution was calculated by summing consecutive frames where the IRAK4/1 signal intensity was above this fluorescent intensity threshold (Fig. 3F, Fig. 4H, Fig. 5E). To calculate the size distribution of MyD88, IRAK4 and IRAK1 assemblies we plotted the max normalized intensity of puncta (Fig. 3D, E, Fig. 4F, G, Fig. 5C, D, Fig. 7F). We restricted this analysis to puncta with a lifetime of 50 seconds or above. We performed data visualization (e.g., of density plots of intensity maxima distribution and percentage of categorized MyD88 puncta per cells) using ggplot2, a data visualization package for R.

## Supplementary Movies

Supplementary Movie 1: The nucleation of multiple MyD88:IRAK4 assemblies on the surface of a live EL4 cell during the 15 minutes after IL-1 stimulation. This movie shows an EL4 cell expressing MyD88-GFP (left panel, green channel in merge) and IRAK4-mScarlet (middle panel, magenta channel in merge) interacting with a SLB functionalized with IL-1β. The movie illustrates the dynamics of MyD88 puncta nucleation and co-assembly with IRAK4 that occurs in response to IL-1. Note the continual nucleation of MyD88 puncta and IRAK4 recruitment over the 15 min movie. Scale bar, 5 µm.

Supplementary Movie 2: The nucleation of multiple MyD88:IRAK1 assemblies on the surface of a live EL4 cell during the 15 minutes after IL-1 stimulation. This movie shows an EL4 cell expressing MyD88-GFP (left panel, green channel in merge) and IRAK1-mScarlet (middle panel, magenta channel in merge) interacting with a SLB functionalized with IL-1β. The movie illustrates the dynamics of MyD88 puncta nucleation, and the downstream recruitment and co-assembly with IRAK1. Note the continual nucleation of MyD88 puncta and IRAK1 recruitment over the 15 min movie. The movie was acquired using TIRF microscopy. Scale bar, 5 µm.

Supplementary Movie 3: Pharmacological inhibition of IRAK4 kinase activity and auto-phosphorylation does not inhibit co-assembly with MyD88. This movie shows EL4 cells endogenously expressing MyD88-GFP and IRAK4-mScarlet interacting with IL-1-functionalized SLBs in presence of DMSO (top row) or 20 µM IRAK4 kinase inhibitor (bottom row). This movie shows that MyD88:IRAK4 co-assembly is undiminished in the presence of IRAK4 kinase inhibition. The movie was acquired using TIRF microscopy. Scale bar, 5 µm.

Supplementary Movie 4: The IRAK4 kinase domain and activity are dispensable for co-assembly with MyD88. A split screen movie showing EL4-MyD88-GFP/IRAK4-KO cells reconstitute with IRAK4-WT-mScarlet (top row), IRAK4^K213/214A^-mScarlet (middle row) or IRAK4^DD^-mScarlet (bottom row) interacting with IL-1-functionalized SLBs. All movies shown acquired using TIRF microscopy. In the merge panels IRAK4 alleles are shown in magenta and MyD88-GFP in green. This movie shows that all WT and kinase domain IRAK4 mutants have the same recruitment and assembly dynamics with MyD88-GFP puncta. We conclude that the IRAK4 death domain is solely responsible for IRAK4 assembly with MyD88. Scale bar, 5 µm.

Supplementary Movie 5: IRAK4 kinase inhibitor reduces the recruitment and incorporation of IRAK1 in nascent Myddosomes. A split screen movie showing EL4-MyD88-GFP/IRAK1-mScarlet cells treated with DMSO (top row) or 20 µM of kinase inhibitor (bottom row) interacting with IL-1-functionalized SLBs. All movies shown acquired using TIRF microscopy. In the merge panels IRAK1-mScarlet images are shown in magenta and MyD88-GFP in green. This movie shows that inhibiting IRAK4 kinase activity reduces the recruitment and stable association of IRAK1 at assembling Myddosome. Scale bar, 5 µm.

Supplementary Movie 6: IRAK4:IRAK1 co assembly required IRAK4 kinase activity. Split screen showing EL4-3xKO cells reconstituted with MyD88, IRAK4^DD^-GFP (Top) or IRAK4^K213/14A^-GFP (Bottom) and IRAK1-mScarlet interacting with IL-1-functionalized SLBs. All movies shown acquired using TIRF microscopy. In the merge panels IRAK2 images are shown in magenta and MyD88-GFP in green. Scale bar, 5 µm.

Supplementary Movie 7: IRAK4 autophosphorlyation potentiates the recruitment and incorporation of IRAK2 into nascent Myddosomes. Split screen showing EL4-MyD88-GFP/IRAK1-KO cells expressing IRAK2-mScarlet treated with DMSO (top row) or 20 µM of IRAK4 kinase inhibitors interacting with IL-1-functionalized SLBs. All movies shown acquired using TIRF microscopy. In the merge panels IRAK2 images are shown in magenta and MyD88-GFP in green. Scale bar, 5 µm.

Supplementary Movie 8: IRAK4 autophosphorlyation potentiates the recruitment and incorporation of IRAK3 into nascent Myddosomes. Split screen showing EL4-MyD88-GFP/IRAK1-KO cells expressing IRAK3-mScarlet treated with DMSO (top row) or 20 µM of IRAK4 kinase inhibitors interacting with IL-1-functionalized SLBs. All movies shown acquired using TIRF microscopy. In the merge panels IRAK3 images are shown in magenta and MyD88-GFP in green. Scale bar, 5 µm.

**Supplementary Figure S1.**
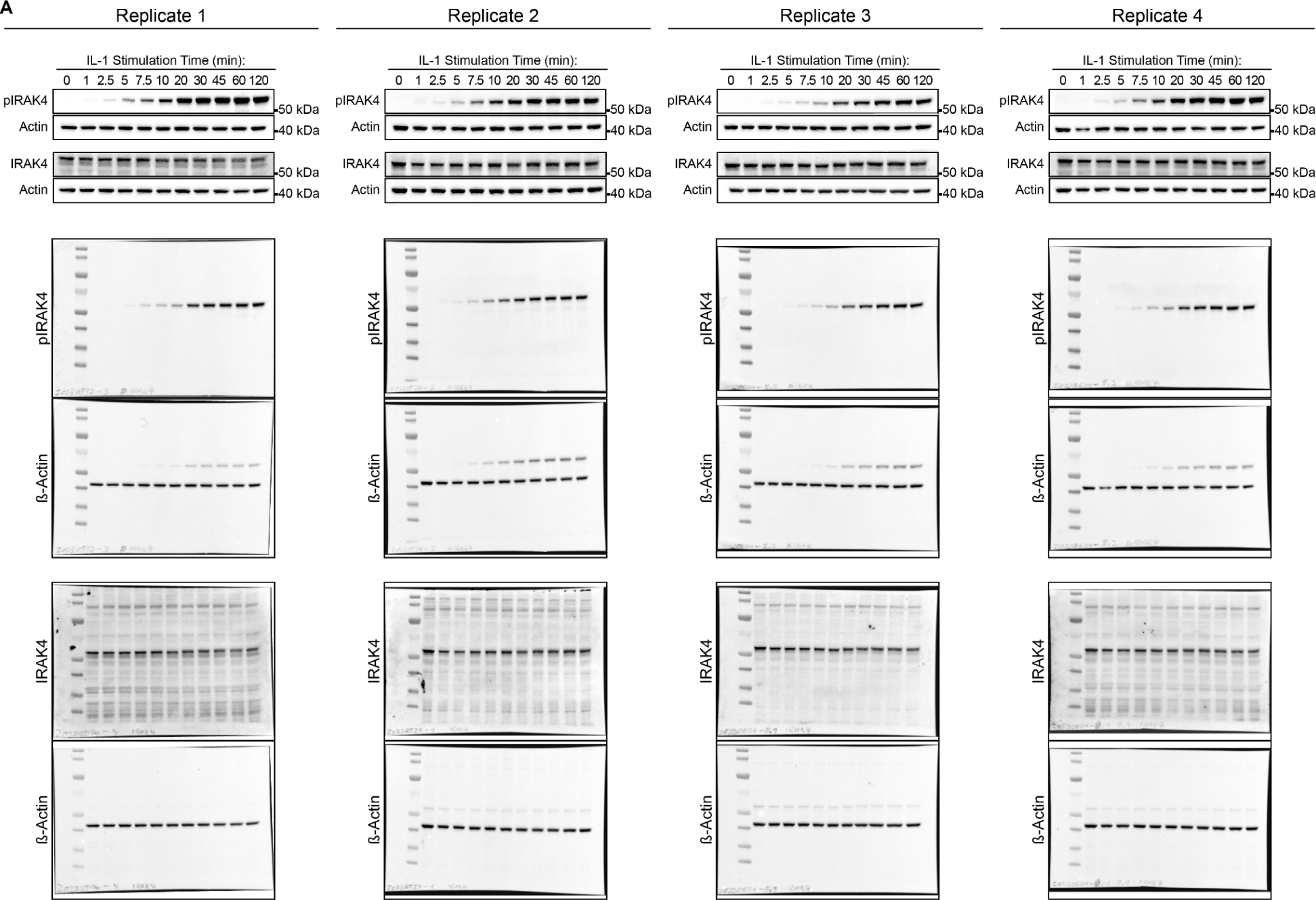
Western blot replicates of the time course analysis of pIRAK4 production after IL-1 stimulation. The full length blot is shown below the cropped blot. Replicate 3 is shown in Fig. 1E. The analysis of each replicate was used to generate the time course of pIRAK4 production in Fig. 1F.

**Figure S2.**
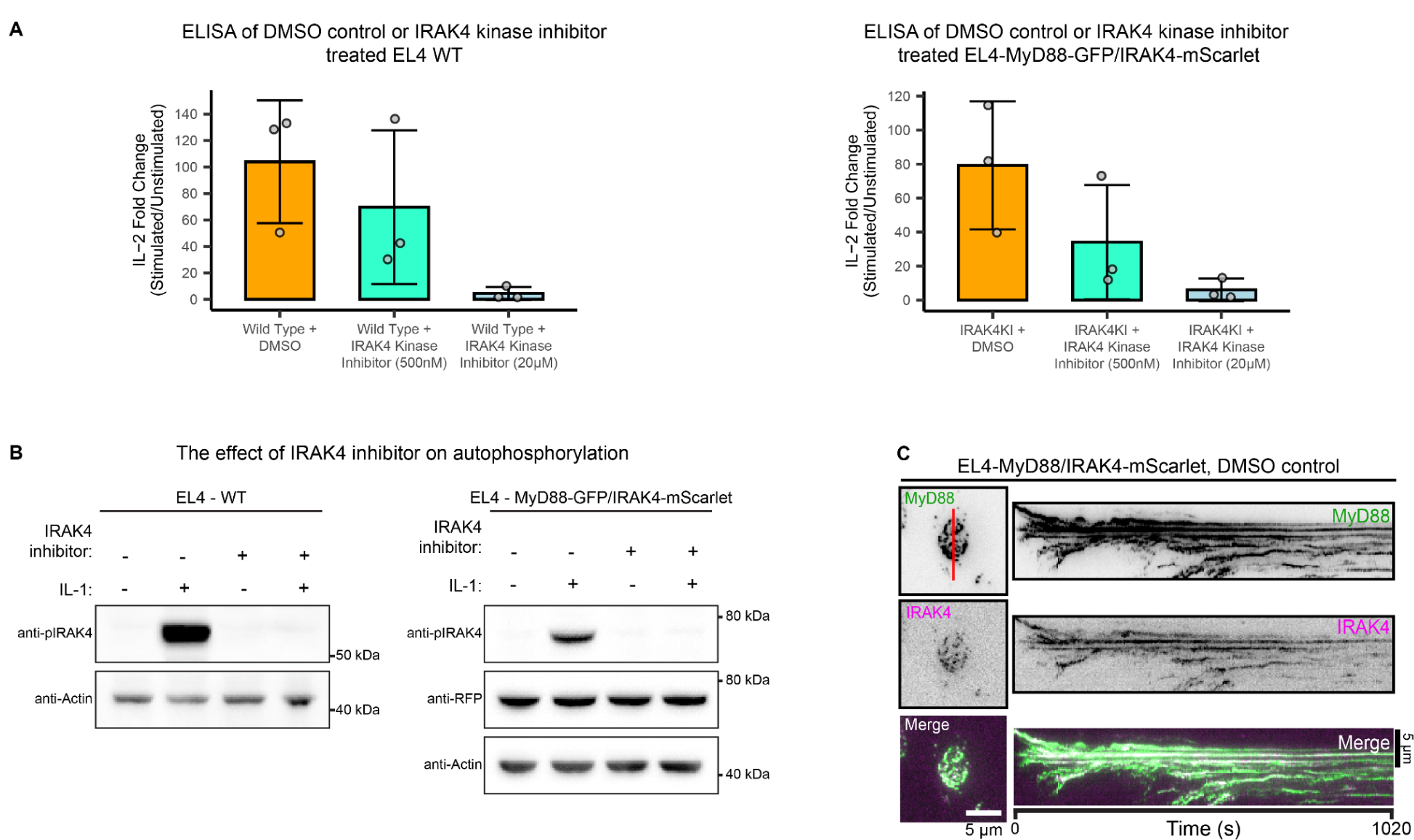
Inhibition of IRAK4 kinase activity suppresses downstream signaling and pIRAK4 production in EL4 cells. **a)** IL-2 release in WT and EL4-MyD88-GFP/IRAK4-mScarlet cells treated with DMSO or IRAK4 kinase inhibitor. IL-2 release was measured by ELISA 24 h after IL-1β stimulation. Values shown are the fold change in IL-2 release. Average values calculated from three independent experiments. Bars represent mean ± SEM. **b)** Treatment with the IRAK4 kinase inhibitor blocks pIRAK4 production after IL-1 stimulation. WT and EL4-MyD88-GFP/IRAK4-mScarlet cell lysates analyzed for pIRAK4 production by Western blot analysis. Cells treated with 20 µM of IRAK4 kinase inhibitor (or DMSO) for 4 h before being stimulated with 1 ng/ml of IL-1β for 30 mins in the presence of the inhibitor. **c)** TIRF images and kymograph analysis of DMSO control treated EL4-MyD88-GFP/IRAK4-mScarlet stimulated on IL-1 functionalized SLBs. Kymographs derived from red line overlaid TIRF images (left panel). Scale bar, 5 µm.

**Figure S3.**
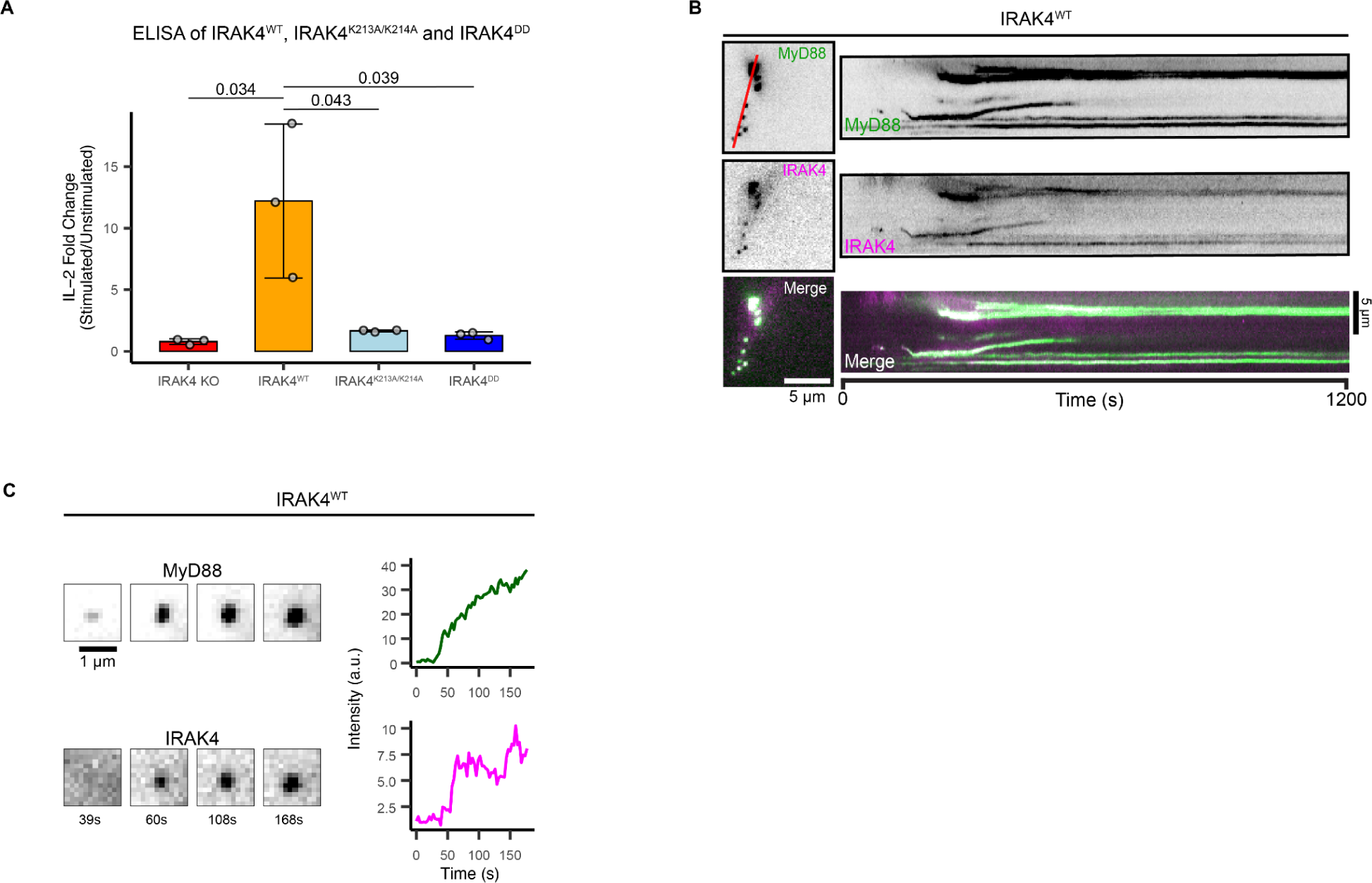
Reconstitution of IRAK4 KO cells with IRAK4^K213/214A^ and IRAK4^DD^ fails to rescue downstream signaling. **a)** IL-2 release in EL4-MyD88-GFP/IRAK4-KO cells reconstituted with IRAK4^WT^, IRAK4^K213/14A^ and IRAK4^DD^. IL-2 release was measured by ELISA 24 h after IL-1β stimulation. Values shown are the fold change in IL-2 release. Average values calculated from three independent experiments. Bars represent mean ± SEM. One way Anova was used to compare the replicate means. **b)** TIRF images and kymograph of EL4-MyD88-GFP/IRAK4-KO cells reconstituted with IRAK4^WT^-mScarlet stimulated on IL-1 functionalized SLBs. Kymographs derived from red line overlaid TIRF images (left panel). Scale bar, 5 µm. **c)** Time-series TIRF images and fluorescence-intensity time series showing the formation of a single MyD88-GFP:IRAK4^WT^ assembly in EL4-MyD88-GFP/IRAK4-KO cells reconstituted with IRAK4^WT^-mScarlet. Scale bar, 1 µm.

**Figure S4.**
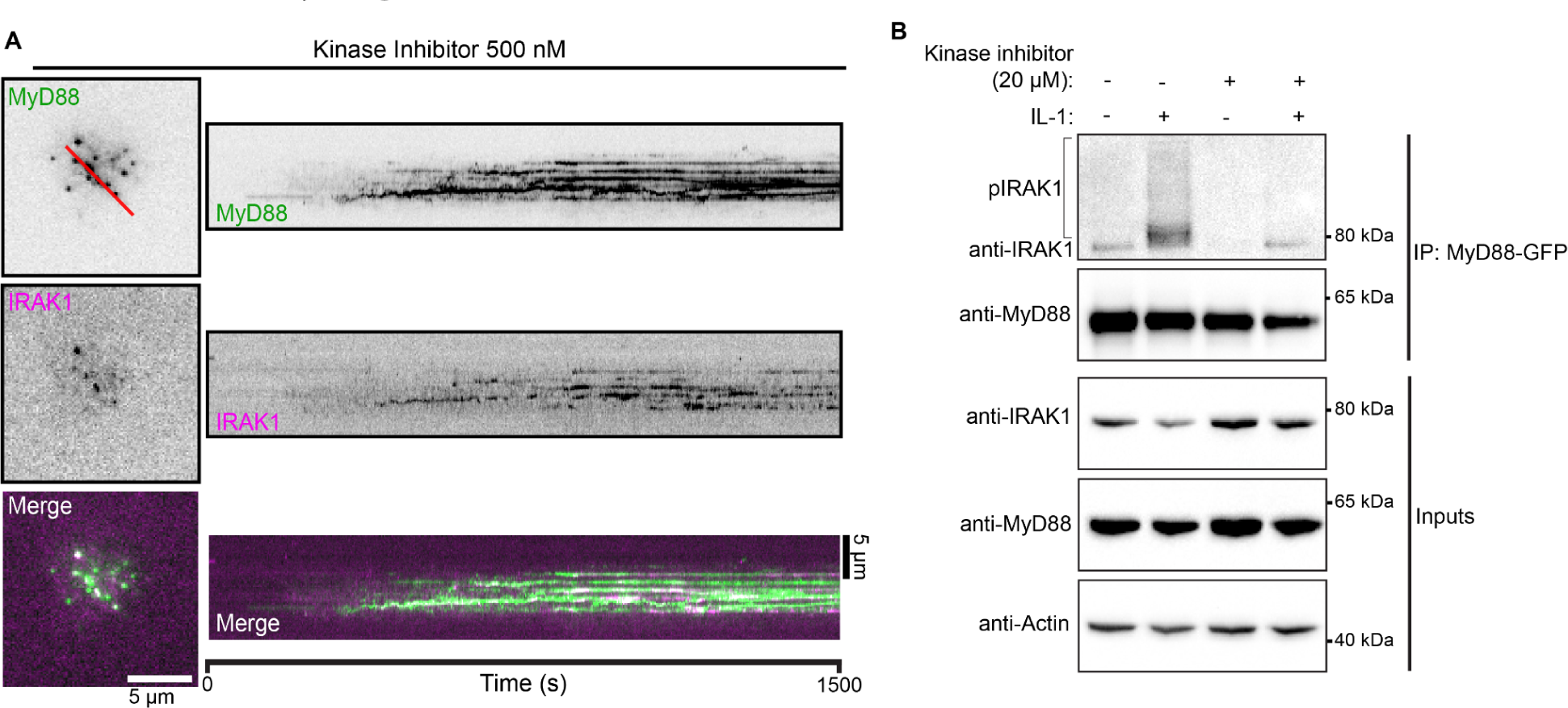
Inhibition of IRAK4 kinase activity disrupts IRAK1 incorporation in Myddosome and downstream post-translational modifications of IRAK1. **a)** TIRF microscopy images and kymograph analysis of IRAK1 recruitment to MyD88 in the presence of 500 nM kinase inhibitor. Scale bar, 5 µm. **b)** Additional Replicate of Myddosome immunoprecipitation from EL4-MyD88-GFP cells. MyD88-GFP was immunoprecipitated from cell lysates treated without or with IRAK4 kinase inhibitor and analyzed by western blot. Treatment with IRAK4 kinase inhibitor blocks the post-translational modification of IRAK1, note the lack of pIRAK1 in lysates from cells treated with kinase inhibitor.

**Figure S5.**
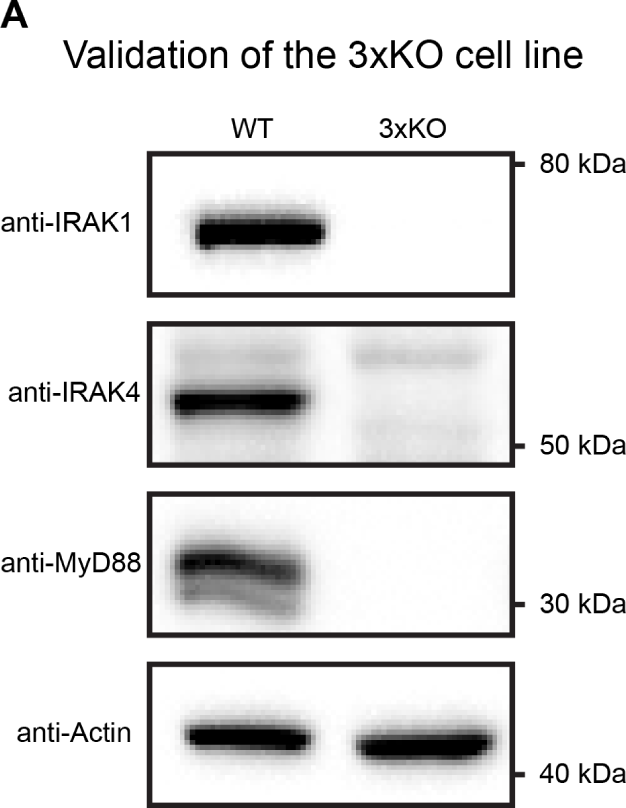
Western blot validation of a EL4 cell clone that is triple KO for MyD88, IRAK4 and IRAK1.

**Figure S6.**
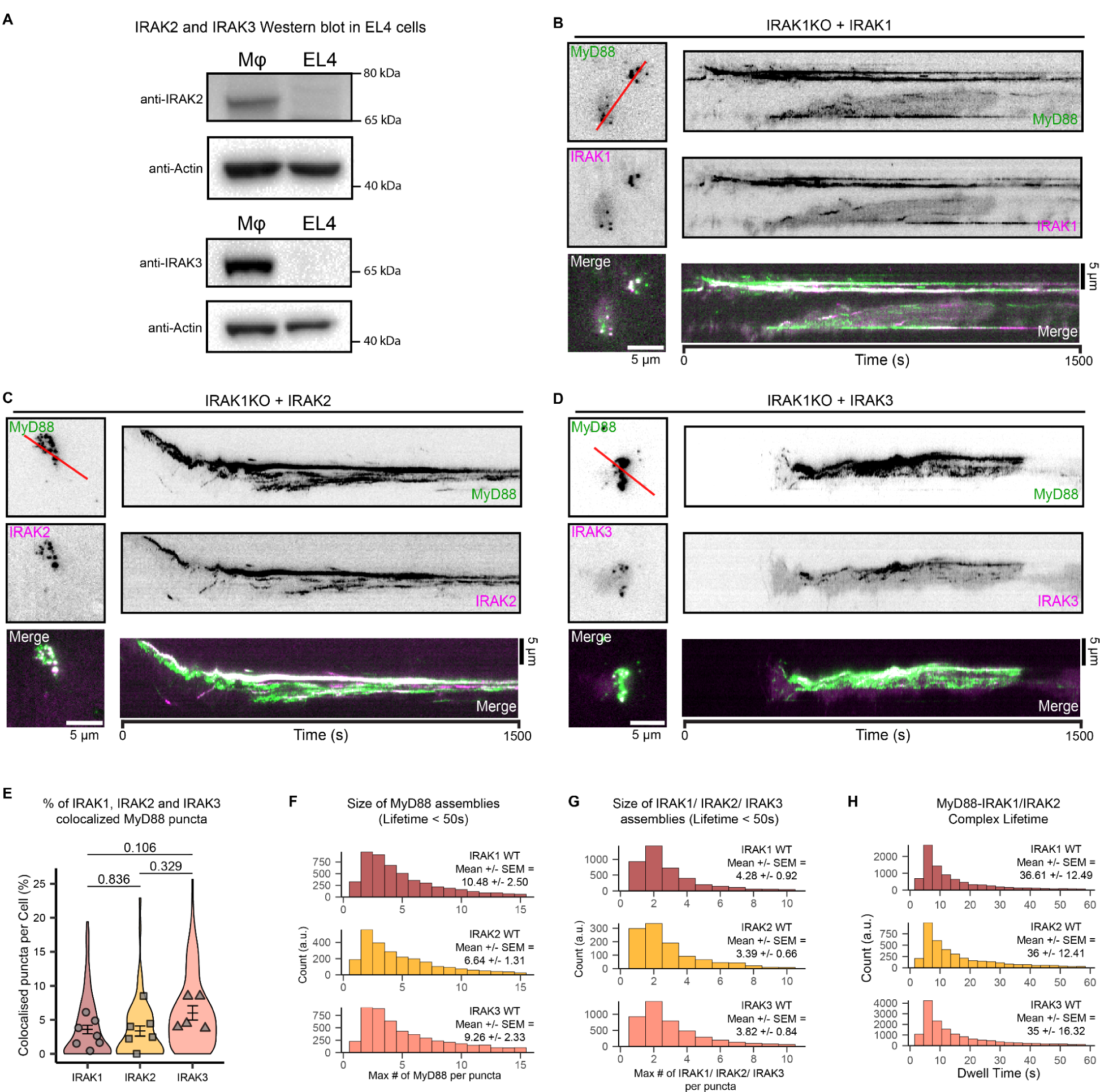
IRAK1-3 have equivalent recruitment and assembly kinetics with MyD88 in reconstituted IRAK1 KO cells. a) Western blot analysis of IRAK2 and IRAK3 expression in EL4 cells. Lysates from bone derived mouse macrophages included as a positive control. **b-d)** TIRF images and kymograph analysis of EL4-MyD88-GFP/IRAK1-KO cells reconstituted with IRAK1-mScarlet (b), IRAK2-mScarlet (c) IRAK3-mScarlet (d) stimulated on IL-1 functionalized SLBs. Kymographs derived from red line overlaid TIRF images (left panel). Scale bar, 5 µm. **e)** The probability of recruitment and co-assembly of IRAK1/2/3 with MyD88 is equivalent and shows no statistical difference. Percentage of MyD88 puncta that colocalize with IRAK1, IRAK2 or IRAK3 per cell in reconstitute EL4-MyD88-GFP/IRAK1-KO cells. Violin plots show the distribution of individual cell measurements. The dots superimposed on the violin plots represent the mean value of each replicate (n = 7, 6 and 5 replicates for IRAK1, IRAK2 and IRAK3 respectively, each replicate encompasses measurements from 8-37, 1-26 and 15-32 cells for IRAK1, IRAK2 and IRAK3 respectively). Bars represent the mean and SEM. Wilcoxon Ranked sum test was used to compare the replicate means. **f-g)** The size of MyD88 and IRAK1/2/3 assemblies are equivalent in reconstituted EL4-MyD88-GFP/IRAK1- KO cells. Histogram plots showing the size distribution of (f) MyD88 assemblies (n = 5961, 2708 and 5879 puncta for IRAK1, IRAK2 and IRAK3 respectively) and (g) IRAK1/2/3 assemblies (n = 4429, 1155 and 4705 puncta for IRAK1, IRAK2 and IRAK3 respectively) in EL4-MyD88-GFP/IRAK1-KO cells reconstituted with IRAK1 (upper, red), IRAK2 (middle, yellow) and IRAK3 (lower, pink). In panel (f), histograms are composed of n = 5961, 2708 and 5879 MyD88 assemblies from cells expressing IRAK1, IRAK2 and IRAK3 respectively. In panel (g), histograms are composed of n = 4429, 1155 and 4705 IRAK1/2/3 assemblies respectively. **h)** Histogram shows the distribution MyD88:IRAK1/2/3 complex lifetimes (seconds) in EL4-MyD88-GFP/IRAK1- KO cells (N = 10057, 4356 and 16912 MyD88:IRAK1/2/3 complexes respectively), from a single replicate.

**Figure S7.**
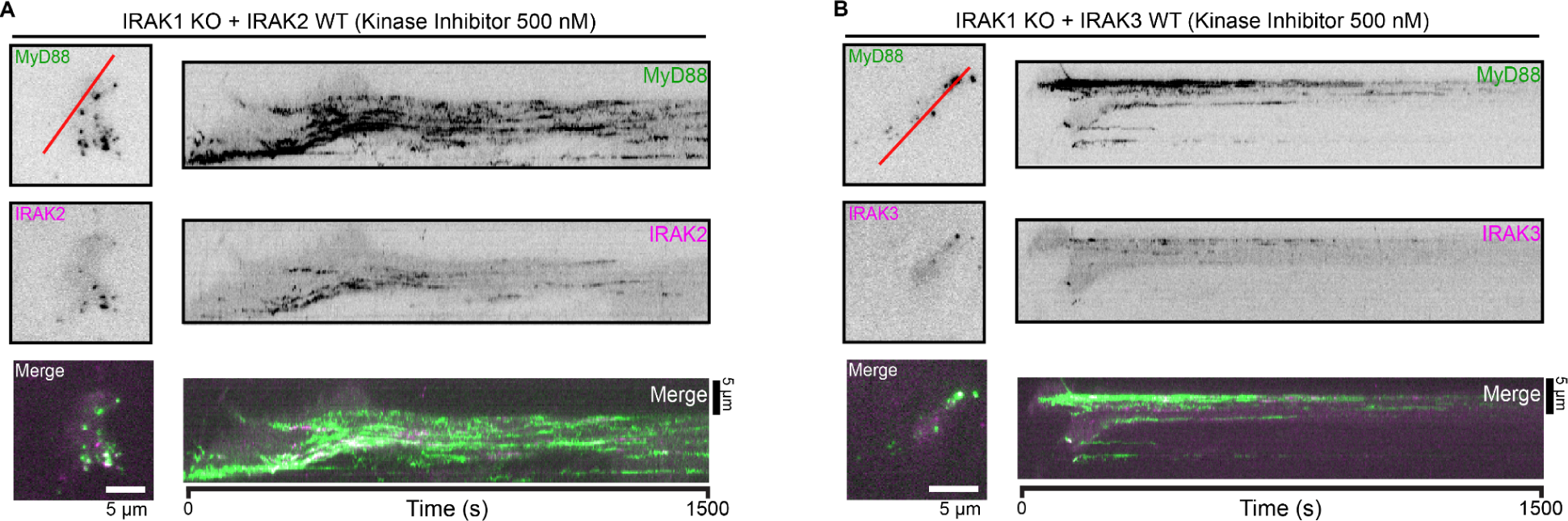
IRAK2/3 recruitment in the presence of 500 nM of IRAK4 kinase inhibitor **a-b)** TIRF images and kymograph analysis of 500 nM IRAK4 kinase inhibitor treated EL4-MyD88-GFP/IRAK1-KO reconstituted with IRAK2-mScarlet (a) or IRAK3-mScarlet (b). Kymographs derived from red line overlaid TIRF images. Scale bar, 5 µm.

**Table 1.**
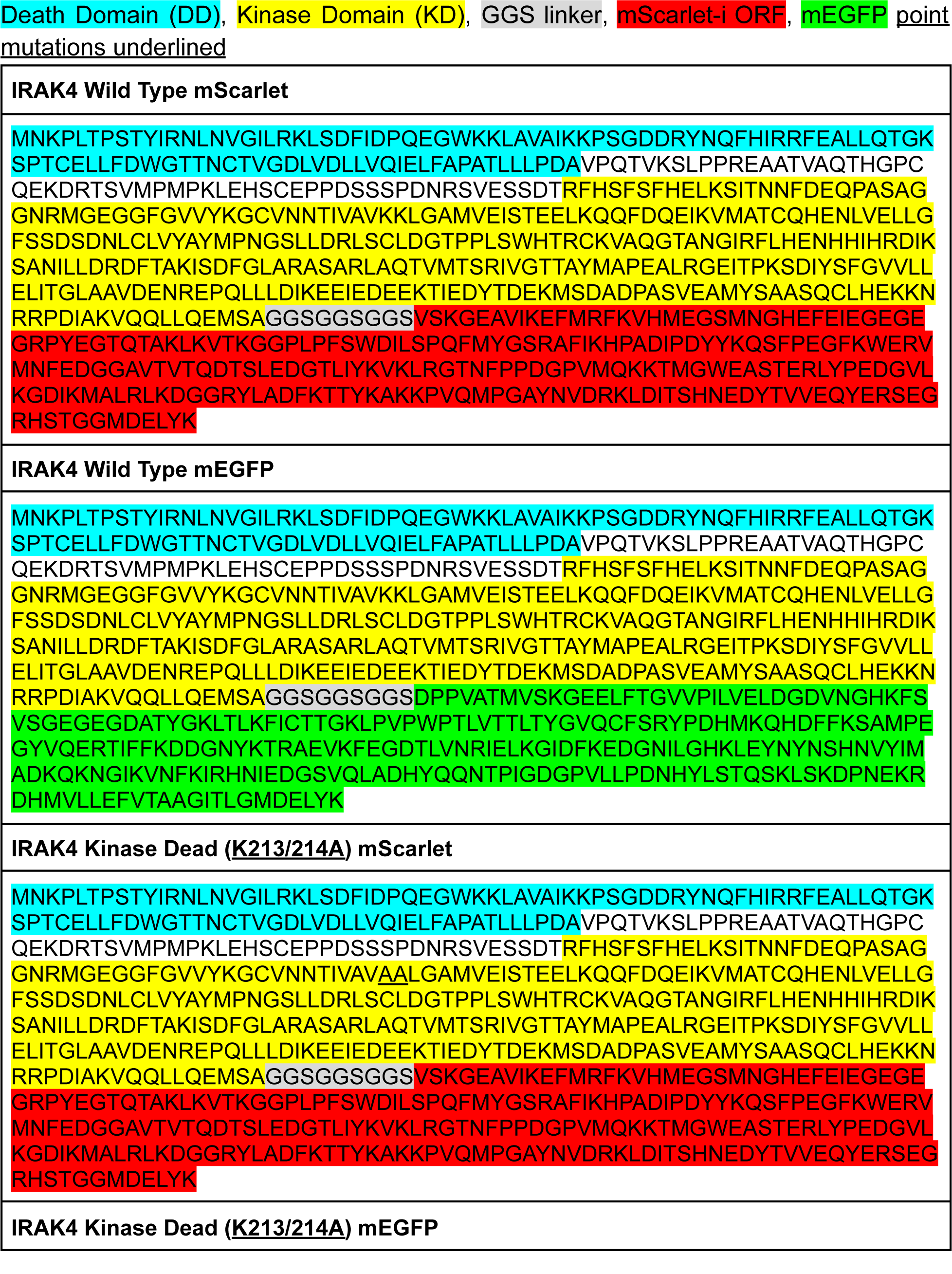
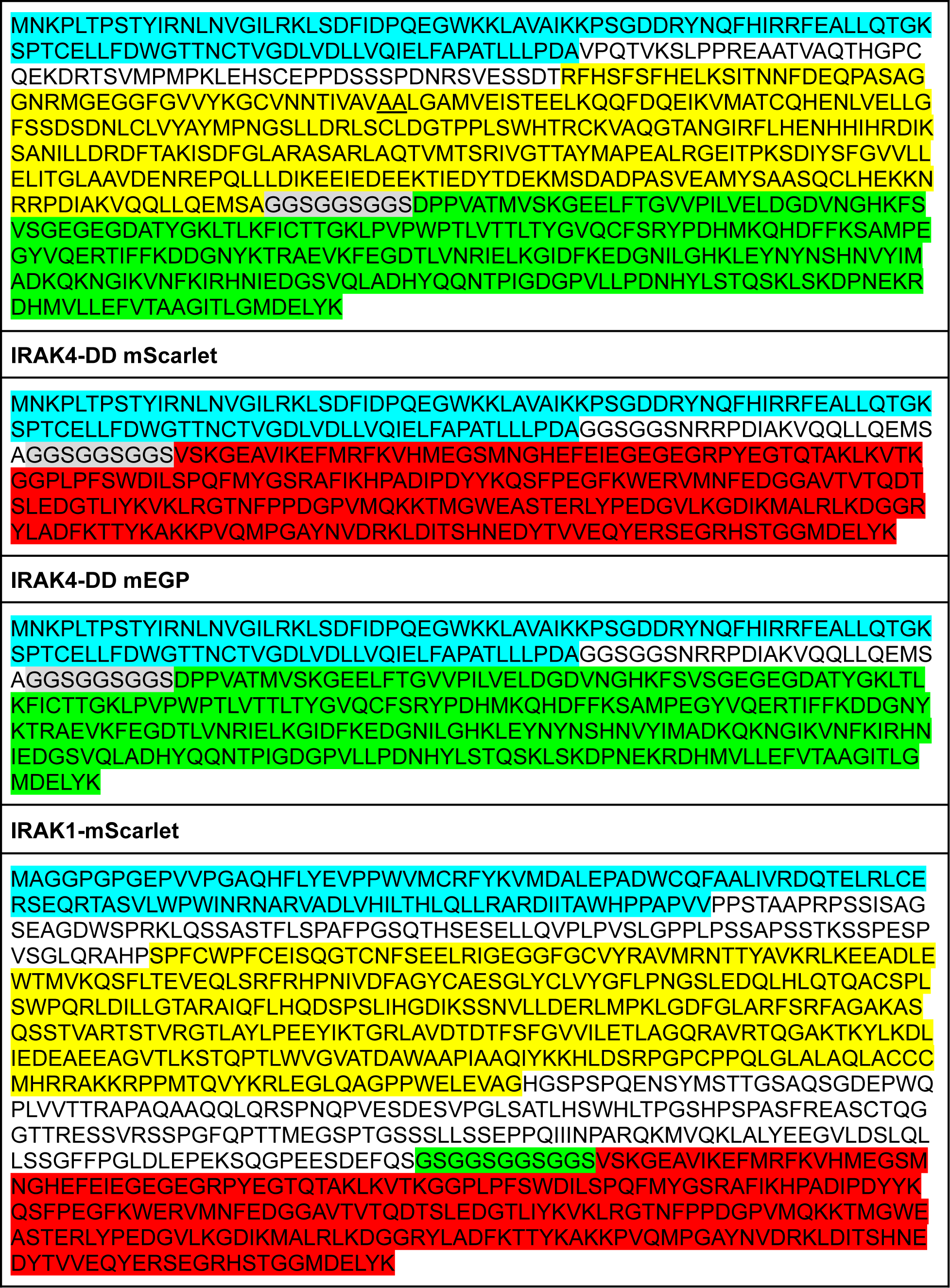
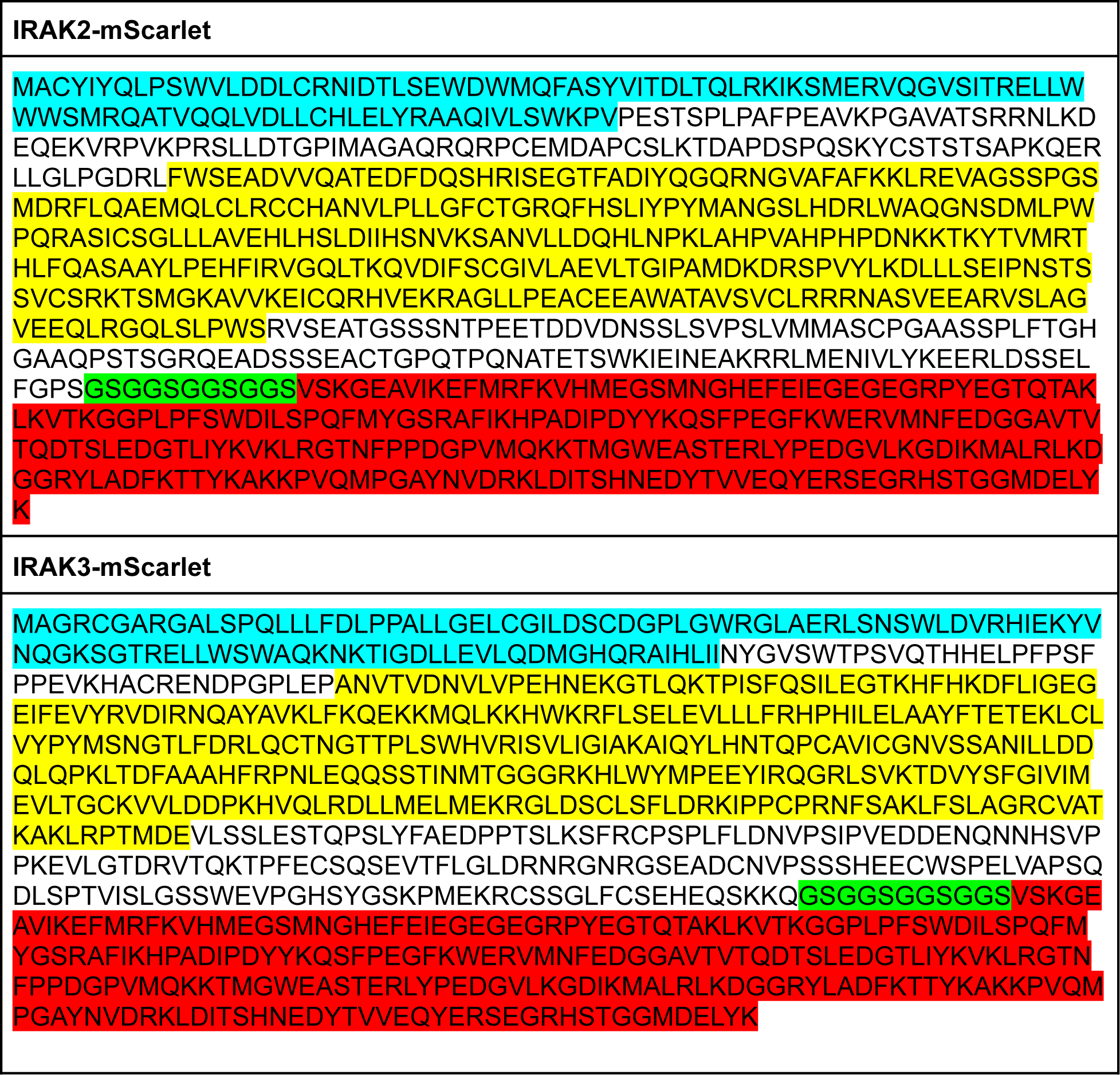
Protein sequences of IRAK1/3/4 mScarlet fusion constructs.

